# EpiCycle: A flexible approach for scalable, three-dimensional reconstructions of complex epithelial tissues

**DOI:** 10.64898/2026.11.20.767655

**Authors:** Elgene J. A. Quitevis, Erik C. Black, Eric Peterman, Louise Harari, Denis J. Headon, James D. Glover, Jeffrey P. Rasmussen

## Abstract

Understanding organ development and repair requires methods to precisely describe cell geometry. Recent image analysis methods allow increasingly accurate segmentation of simple epithelial tissues. However, many vertebrate organs contain more complex stratified epithelia, for which accurate 3D cellular segmentation remains challenging. Here, we describe EpiCycle, an iterative, user-friendly approach developed to address the unique segmentation challenges posed by stratified epithelia. Starting from sparse or no ground truth annotations, EpiCycle guides users through a human-in-the-loop refinement process that culminates in 3D consensus reconstruction. Through cyclical correction, annotation and model retraining, EpiCycle allows users to progressively build robust, dataset-specific segmentation models while simultaneously generating 3D ground truth. To highlight the utility of EpiCycle, we train models that generate high-confidence, tissue-level segmentation of the stratified squamous epithelium in adult zebrafish skin. We apply our approach to segmentation of mouse embryonic branching salivary glands demonstrating its versatility across organs and species. Finally, we produce and analyze quantitative measurements of stratified cell geometry in human fetal epidermis. Thus, EpiCycle enables scalable and quantitative 3D reconstructions of complex vertebrate tissues without relying on pre-existing ground truth.

## INTRODUCTION

Organs adopt diverse and complex three-dimensional architectures to perform specific functions. To generate mature organ shapes, constituent cells undergo dynamic shape changes and rearrangements during development (Gómez-Gálvez et al., 2021; Lemke and Nelson, 2021). Damage to adult tissues can trigger rapid changes in cell shape necessary to restore organ architecture (Lim et al., 2024). Thus, methods to precisely describe cell geometry are fundamental to understanding organ development and repair.

Recent advances in fluorescence microscopy and deep learning (DL) image segmentation methods now permit reconstruction of cell morphology from confocal micrographs (Moen et al., 2019), thus allowing description of cell shape in diverse organs across time and perturbations. Robust methods exist for segmentation of simple epithelia, including those that have deformed geometries (Achard et al., 2025; Aigouy et al., 2020; Andrés-San Román et al., 2023; Cilla et al., 2015; Erguvan et al., 2019; Merle et al., 2023; Wang et al., 2022). However, many vertebrate organs—such as skin, cervix, thymus and salivary gland—contain pseudostratified or stratified epithelia. DL-based reconstruction of cell shape in these contexts remains challenging (Gómez et al., 2021), owing to the variable image quality, crowded tissue microenvironments and complex cell shapes.

Widely used DL-based segmentation tools such as Cellpose (Stringer et al., 2021) are often trained on imaging datasets from 2D tissue and cell culture. Application of these tools to complex epithelial organs requires prohibitively dense human-generated annotations to train native 3D segmentation models. An alternative approach to 3D segmentation is to perform 2D segmentation on image orthoviews (*xy*, *xz* and *yz* views) before fusing into 3D. The recently developed Cellpose3D, CellStitch, 3DCellComposer, Bootstrapper and u-Segment3D methods demonstrate the utility of this 2D-to-3D approach, without requiring large training datasets (Chen et al., 2025; Liu et al., 2023; Stringer et al., 2021; Thiyagarajan et al., 2024; Zhou et al., 2025). Yet, these pipelines are typically used as one-shot batch procedures. In practice, segmenting densely layered, anisotropic epithelia with tightly packed cells still requires extensive manual curation and model fine-tuning when using these methods. This motivates approaches that explicitly couple interactive 2D annotation, retraining and 3D consensus reconstruction.

Here, we present EpiCycle, a workflow designed to extend existing segmentation tools to allow iterative, human-in-the-loop (HITL) improvement of 2D-to-3D segmentation. EpiCycle operates as a supervised, user-driven graphical interface in the widely used image viewer napari (Sofroniew et al., 2025) in which users validate model predictions and steer model updates through cyclical refinement loops. We show that EpiCycle enables the development of customizable models that adapt to tissue-specific structures, imaging conditions and scientific priorities defined by the user, and offers a unified platform that enables scalable 3D reconstructions and quantitative studies of complex epithelia.

## RESULTS

### EpiCycle: An iterative 2D-3D human-in-the-loop segmentation workflow

EpiCycle is a two-phase HITL workflow that integrates iterative 2D model refinement with 3D reconstruction. In Phase 1, the user selects representative stacks and annotates a small number of lateral (*xy*) and/or axial (*xz/yz*) slices (**Fig. 1, Phase 1**). EpiCycle uses these curated mask annotations to train an initial pair of Cellpose-based lateral and/or axial models, which are then reapplied to the same stacks to generate new predictions. Users correct errors, append the curated slices to the training set, and retrain the model, forming an iterative cycle that incrementally expands the labeled dataset and simultaneously improves 2D segmentation performance of the model at each iteration. This cyclical process follows the general HITL paradigm in tools such as Cellpose, in which user-corrected 2D examples iteratively refine model performance. EpiCycle extends this paradigm to coordinate lateral and axial model training to support multi-orthoview consistency required for downstream 3D reconstruction.

**Figure 1.**
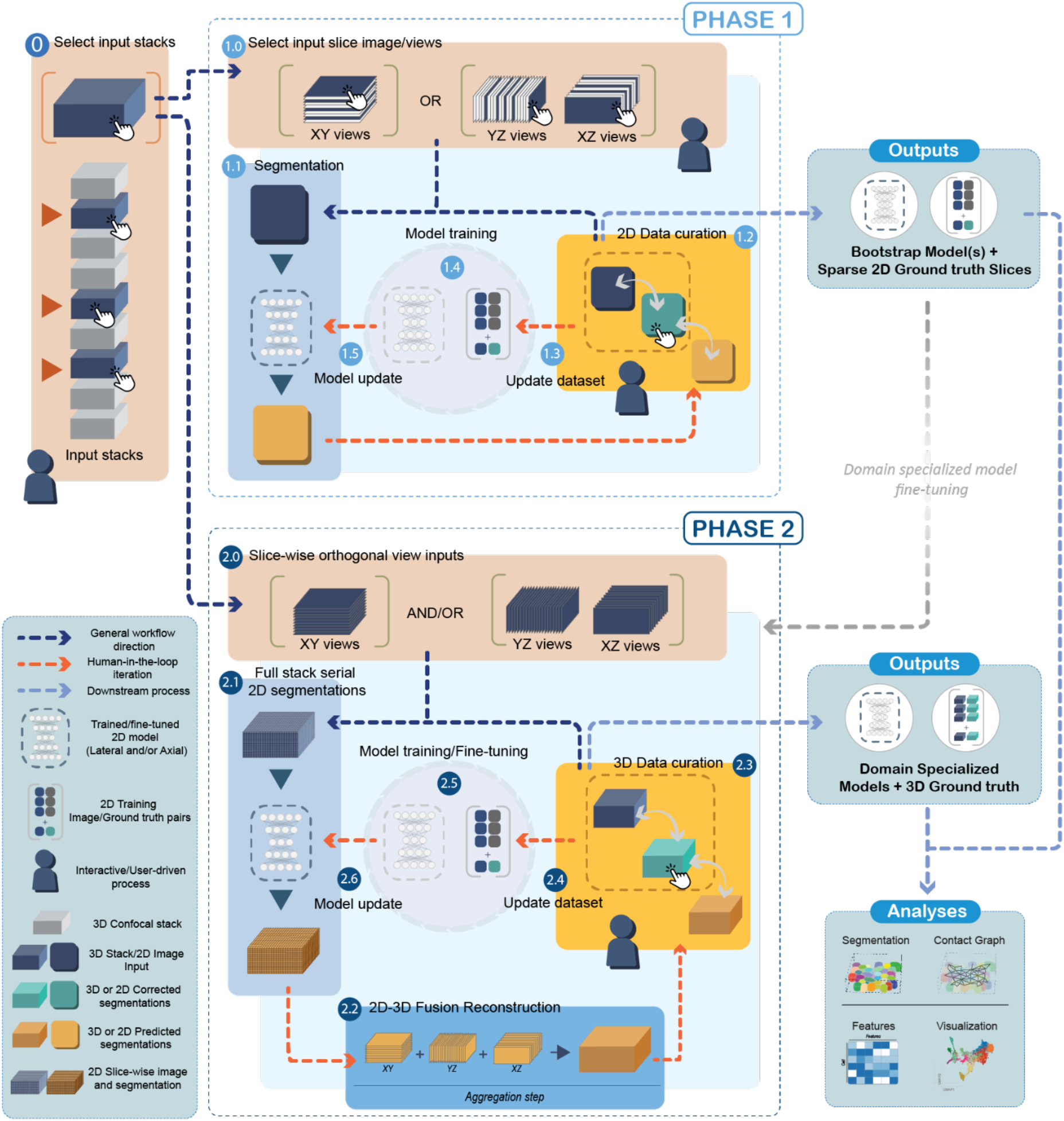
EpiCycle: an iterative 2D-3D human-in-the-loop segmentation approach. EpiCycle operates in two phases that together define an annotation-training cycle in which curated 2D examples drive progressive improvement of segmentation models supporting 3D reconstruction. **Phase 1 (top):** iterative 2D model refinement via human-in-the-loop workflow. (0) The user selects one or more representative 3D image stacks from the input dataset. (1.0) Sparse 2D slice views are extracted in one or more orthogonal orientations (*xy* lateral; *xz*, *yz* axial). (1.1) An initial Cellpose-based lateral and/or axial model pair generates 2D segmentation predictions on selected slices. (1.2) The user reviews predictions and curates corrected 2D annotations interactively. (1.3) Curated 2D image-annotation pairs are appended to the cumulative training dataset. (1.4) The lateral and/or axial models are retrained on the updated dataset. (1.5) Updated models replace the previous models and regenerate predictions, beginning the next iteration of the cycle. Phase 1 outputs a set of bootstrap model(s) and accumulated sparse 2D ground truth slices. **Phase 2 (bottom):** 3D reconstruction and model refinement. (2.0) Bootstrap models are applied serially across full 3D stacks in all three orthogonal views (*xy, xz, yz*). (2.1) This generates dense 2D segmentation predictions for each orientation (probability maps and flow predictions). (2.2) Orthogonal 2D predictions are fused into volumetric instance segmentations using the direct method in u-Segment3D (Zhou et al., 2025). (2.3) The user reviews 3D reconstructions and curates corrections at the volumetric level. (2.4) Corrected 3D segmentations contribute to an updated training dataset that may include additional 2D slice annotations derived from the corrected volumes. (2.5) Models are fine-tuned or retrained from scratch using the expanded curated dataset. (2.6) The updated domain specialized models replace earlier models, and a new cycle begins and previously non-segmented patches are processed. Phase 2 outputs domain-specialized models and 3D ground truth volumes that support downstream tissue-scale analyses.

In Phase 2, EpiCycle runs the refined lateral and axial models serially across entire stacks to generate dense 2D predictions in all three image orthoviews (**Fig. 1, Phase 2**). These predictions are fused into volumetric segmentations using a slicewise 2D-to-3D reconstruction method implemented in u-Segment3D, after which users may perform optional 3D-level corrections. The user can either directly use the corrected 3D instances for analysis or convert them back into 2D slices to continue improving existing models or to train new models. A finalized model can then be applied to new target stacks or tiled whole-tissue volumes, from which EpiCycle extracts per-cell logical features and cell-cell contact graphs for downstream exploratory analyses. Together, these two phases define an annotation-training cycle in which curated 2D examples drive progressive segmentation model improvement that ultimately supports 3D reconstruction.

### Human-in-the-loop training progressively improves 2D lateral and axial segmentation

As our primary system for developing EpiCycle we chose the adult zebrafish trunk epidermis, an optically accessible stratified squamous epithelium used to study skin development and regeneration (Aman et al., 2018; Aman et al., 2023; Brown et al., 2023; Chen et al., 2016; Craig et al., 2025; Le Guellec et al., 2004; Peterman et al., 2023; Peterman et al., 2024; Rasmussen et al., 2018; Richardson et al., 2016; Roan et al., 2021; Santoso et al., 2024). The trunk epidermis drapes across the outer surface of scales, which are bony plates that form the dermal appendages of zebrafish (Aman and Parichy, 2024; Le Guellec et al., 2004) (**Fig. S1A**). The adult epidermis contains three molecularly distinct layers of epithelial keratinocytes: superficial, suprabasal and basal cells (Hou et al., 2020; Marí-Beffa et al., 1996) (**Fig. S1A**). To visualize keratinocyte morphologies, we imaged *TgBAC(cdh1:cdh1-sfGFP);Tg(actb2:h2b-2x-mScarlet)* fish, in which E-cadherin/Cdh1-sfGFP labels epithelial cell-cell junctions (Colak-Champollion et al., 2019) and H2B-2x-mScarlet labels all nuclei (Peterman et al., 2025) (**Fig. S1B,C**). Imaging this dual reporter at progressive *z*-depth illustrated layer-specific cell morphologies (**Fig. S1D,E**). From confocal imaging of this dual-reporter line, we curated eleven densely annotated 3D ground truth *z*-stacks from adult trunk epidermis (see Methods; imaging parameters in **Table S1**). We designated seven stacks as model development data for Phase 1 EpiCycle training, and four stacks as independent evaluation data for downstream evaluation.

To simulate a HITL workflow, we used the model development data but only exposed a small number of 2D slices to the model at each training iteration (**Fig. 2A**) (see Methods). Specifically, in each training round, we randomly selected 15 lateral (*xy*) and 30 axial (*xz* and *yz*) slices per stack without replacement, resulting in a cumulative training set that grew linearly before saturating at 700 lateral and 1,400 axial slices by rounds 6-7 (**Fig. S2A**). We sampled 2D masks from the 3D ground-truth annotations and trained separate lateral and axial Cellpose models on the cumulatively expanded training dataset (see Methods). We performed ten rounds of training, scoring each model by average precision at 50% intersection over union (AP@50; **Fig. 2B**), and found that predictions sharpened over the course of training. Models trained after four and nine rounds captured 2D cell boundaries more accurately and reduced under-segmentation and spurious splits in both lateral and axial views compared to the baseline model (round 0) (**Figs. 2C, S2B,C**).

**Figure 2.**
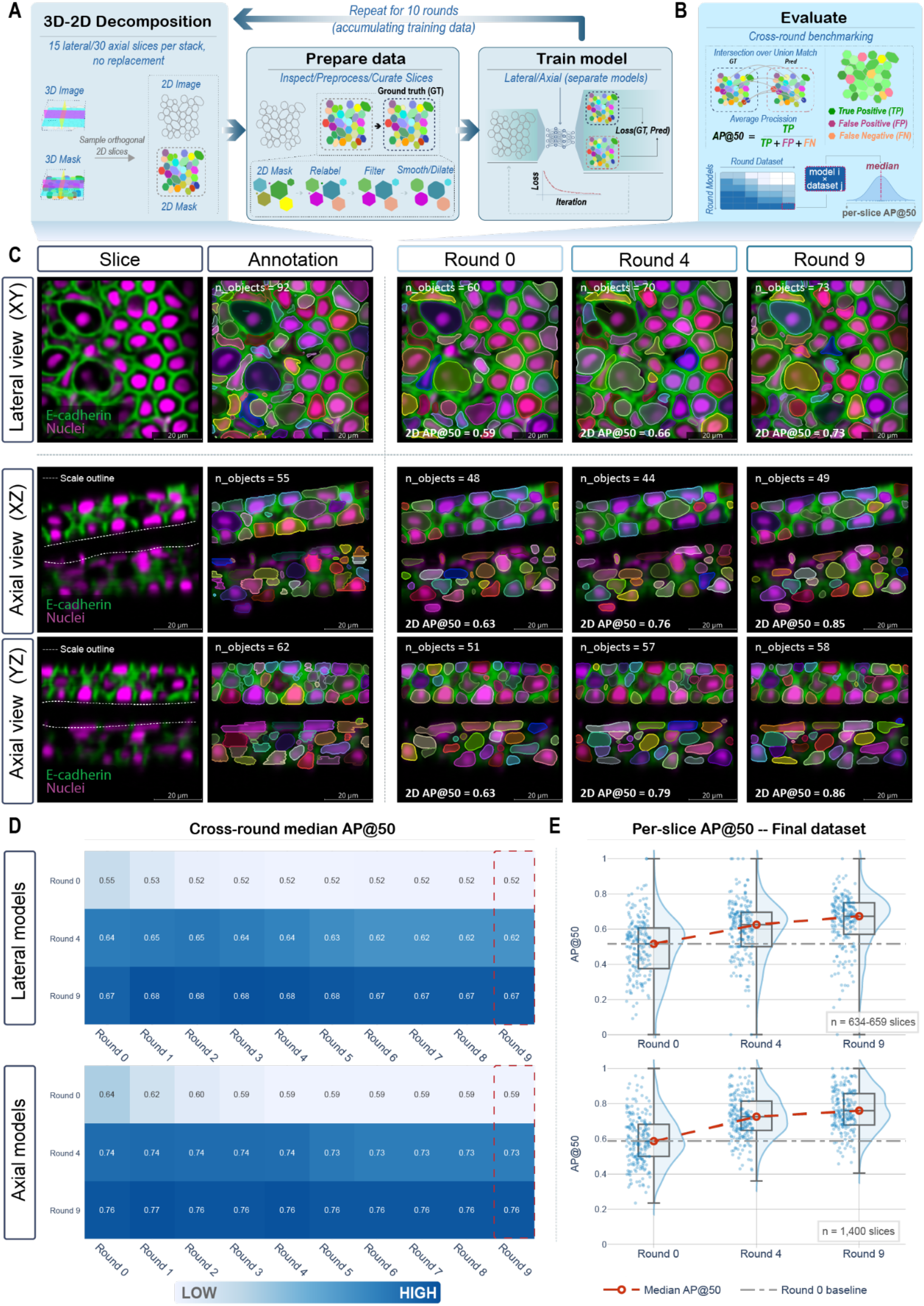
Human-in-the-loop training progressively improves 2D lateral and axial segmentation. **(A)** Simulated HITL bootstrapping workflow. Seven densely annotated 3D ground truth stacks were decomposed into 2D slices (15 lateral and 30 axial slices per stack per round, and curated (2D mask extraction, relabeling, size filtering and morphological smoothing/dilation) before being added to a cumulative training set. Separate lateral and axial models were retrained on the expanding dataset. The cycle was repeated for ten rounds. **(B)** Cross-round benchmarking setup. (Top) Model performance was quantified using average precision at 50% intersection-over-union (AP@50), computed per slice based on matched ground truth (GT) and predicted (Pred) instances. True positives, false positives and false negatives were defined per the IoU threshold. (Bottom) Schematic of the model x dataset performance matrix in which each entry is the median distribution of AP@50. **(C)** Representative 2D confocal slice image, GT image pair and outputs across training rounds. Top: lateral (*xy*) view, Middle and bottom: axial (*xz, yz*) views. Columns from left to right: raw image slice, ground truth annotation, round 0 (baseline), round 4 and round 9 model predictions. **(D)** Cross-round median AP@50 heatmaps for lateral (top) and axial (bottom) models. Rows indicate the training round of the model (rounds 0, 4, 9 shown). Columns indicate the dataset round (rounds 0-9). Dashed red boxes highlight performance on the final round 9 dataset. **(E)** Per-slice AP@50 distributions evaluated on the final round 9 dataset for lateral models (top, *n* = 634-659 slices after excluding empty slices) and axial models (bottom, *n* = 1,400) at rounds 0, 4 and 9. Box plots show median and interquartile ranges with overlaid jittered points and kernel density estimates. Red line: median AP@50 trajectory across rounds. Gray dashed line: round 0 baseline. Per-round medians and slice counts in Table S2.

Next, we quantified how HITL training affected 2D performance over rounds. Cross-round evaluation, in which each model was evaluated on datasets containing annotation from different rounds, showed a consistent increase in median average precision at 50% intersection over union (AP@50), a measure of model performance, for both lateral and axial models (**Fig. 2D**). For the lateral model, median AP@50 along the diagonal (each model evaluated on the dataset available for that round) increased from 0.55 at round 0 to 0.67 at round 9, with most of the gain achieved by rounds 4-6 and values largely saturating thereafter (**Fig. 2D**). When we evaluated all lateral models on the final dataset (dataset round 9), which contained the highest diversity of annotations, median AP@50 increased from 0.52 for the baseline model to 0.67 for the final model (**Fig. 2D,E**).

By contrast, the axial model started from a higher baseline and improved more rapidly. The diagonal median AP@50 increased from 0.64 at round 0 to 0.76 by round 9, with performance reaching 0.73-0.74 by round 3-4 and then plateauing around 0.75-0.76 in later rounds (**Fig. 2D**). On the final dataset, median AP@50 rose from 0.59 for the baseline axial model to 0.76 for the final model (**Fig. 2D,E**). Rows for the later axial models were nearly uniform (0.75-0.76) across all dataset rounds (**Fig. 2D**), indicating that HITL training yields axial segmentation models that perform robustly across the full range of images. Together, these results show that a modest number of curated slices per round is sufficient to train Cellpose-based lateral and axial models that provide reliable 2D segmentation for this dataset context.

### Iterative 2D refinement enables accurate 3D reconstruction

To determine whether improvements in 2D lateral and axial segmentations translate to accurate volumetric cell reconstructions, we applied models from select HITL rounds (0, 4, 9) to full 3D image stacks and reconstructed cell instances using the direct method implemented in u-Segment3D (**Fig. 3A,B**). Briefly, this approach integrates complementary Cellpose foreground-background probability predictions and flow predictions from orthogonal 2D views for 3D instance reconstructions (see Methods). To quantify reconstruction accuracy, we computed 3D AP@50 on the generated 3D masks (see Methods). We observed that reconstruction performance improved modestly across progressive HITL rounds and stacks, with median AP@50 increasing from 0.52 to 0.76 for the round 0 and 9 models, respectively (**Fig. 3C; Table S2**). Qualitatively, reconstructed cells exhibited relatively cleaner boundaries and more coherent cell volumes consistent with sharpening of 2D predictions as previously observed during progressive rounds of HITL training (**Fig. 3D, development stacks**). Notably, we also observed substantial differences in accuracies within the independent evaluation data consistent across different model rounds (**Fig. 3D, test stacks**).

**Figure 3.**
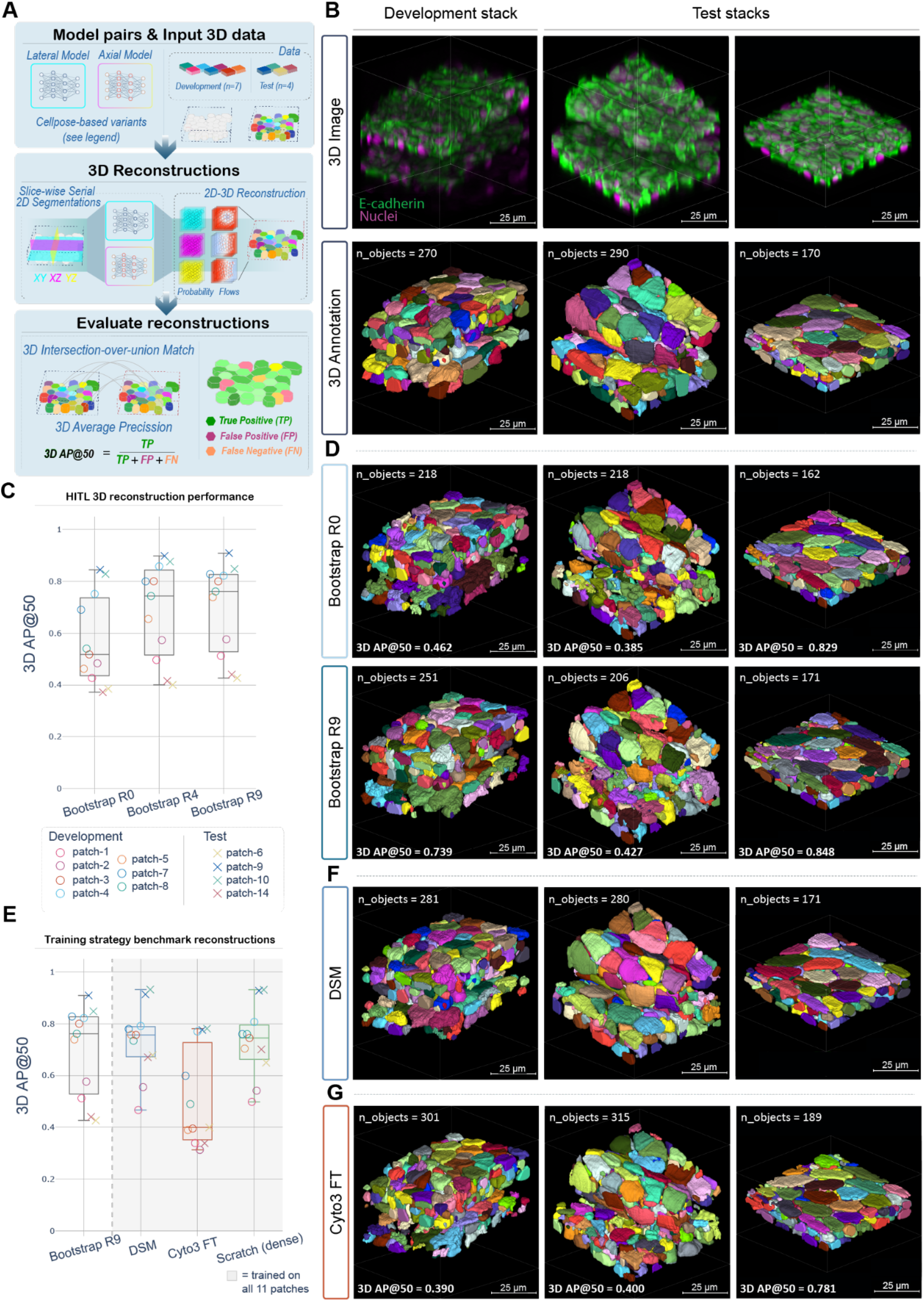
Iterative 2D refinement enables accurate 3D reconstruction of stratified epithelium. **(A)** Schematic of 3D reconstruction and evaluation pipeline. Trained 2D Cellpose model pairs (lateral and axial; see Methods) are applied to full 3D image stacks. Slice-wise 2D segmentations are computed in three orthogonal orientations (*xy, xz, yz*) and combined into 3D instance segmentations via the direct method in u-Segment3D (Zhou et al., 2025), which fuses probability and gradient flow predictions across orthogonal views. Reconstructions are evaluated against 3D annotations using 3D intersection-over-union matching and average precision at 50% IoU threshold. **(B)** Representative 3D image and ground truth annotation for development stack (left column) and two test stacks (right columns). Composite images show cell membranes and nuclei, using the *TgBAC(cdh1:cdh1-sfGFP);Tg(actb2:h2b-2x-mScarlet)* transgenic line. Eleven densely annotated stacks were used in total (7 development, 4 test stacks for independent evaluation; see A). **(C)** Per-stack 3D AP@50 across HITL bootstrap rounds. Bootstrap models (R0, R4, R9) were trained on iteratively accumulated sparse 2D annotations (round 0 from scratch, later rounds continuing from the previous round) from 7 development stacks (see Fig. 2). Each model was applied to all 11 stacks; circles indicate development stacks; crosses indicate test stacks. Box plots show median and interquartile range overlaid with per stack computed 3D AP@50. Per-stack values and summary statistics in Table S2. **(D)** 3D reconstructions of Bootstrap R0 (baseline) and Bootstrap R9 models on the same stacks shown in (B). Reconstructions become qualitatively cleaner across HITL rounds, with fewer irregularities and more coherent and plausible cell volumes by R9. **(E)** Per-stack 3D AP@50 across training conditions, all evaluated on the same 11 stacks. **Bootstrap R9**: scratch-trained model from 9 rounds of sparse HITL on 7 development stacks (same model as in C; shown as reference and Fig. 2). **DSM (Domain Specialized Model):** Bootstrap R4 model further fine-tuned on dense annotations from all 11 stacks (see Methods). **Cyto3 FT**: pretrained Cellpose cyto3 model fine-tuned on dense annotations from all 11 stacks (same training condition as DSM). **Scratch (Dense)**: trained from scratch on dense annotations from all 11 stacks, defining the performance ceiling for the Cellpose architecture on this dataset. Gray shading indicates models trained from all 11 stacks; dashed vertical line separate the sparse-HITL reference (Bootstrap R9) from the dense-supervision strategies. **(F)** 3D reconstructions of the DSM on the same stacks shown in (B). Reconstructions show coherent cell volumes and fewer surface irregularities and fragmentation compared to Cyto3 FT and bootstrap HITL model reconstruction results. **(G)** 3D reconstruction of Cyto3 FT model on the same stacks shown in (B). Despite training on dense annotations from all 11 stacks, reconstructions show visible surface irregularities and fragmentations compared to DSM.

We next asked whether additional dense supervision training could reduce stack-to-stack variability while retaining the benefits of HITL-trained models. We selected the round 4 HITL model, corresponding to early performance saturation, and fine-tuned accordingly using the same curated annotations from all eleven stacks (see Methods). We denote this fine-tuned model as a domain-specialized model (DSM). DSM achieved reduced performance variability and improved worst-case reconstruction accuracy relative to sparse HITL training, approaching the performance of models trained from scratch with dense supervision (**Fig. 3E, scratch; 3F, DSM**). We further asked whether pretrained generalist models could achieve comparable performance by similarly fine-tuning using the same data; however, fine-tuned Cellpose cyto3 models consistently underperformed both the DSM and scratch-trained models (median AP@50 = 0.40; **Table S2**), with additional training and hyper parameter optimization failing to reduce the gap (**Figs. 3E,G, S3A**).

Together, these results demonstrate that: (1) sparse HITL annotation is sufficient for 3D-accurate cell reconstruction; (2) reconstruction performance generalizes to unseen tissue without volumetric ground truth annotation; (3) dense, domain-specific training can further reduce performance variability across heterogeneous samples and establish a performance ceiling for the tissue domain; and (4) pretrained Cellpose priors are not readily adapted to the morphological and photometric properties of complex stratified epithelial context, suggesting that training models from scratch is required to obtain accurate 3D reconstruction in this setting. Importantly, corrected 3D reconstructions produced in EpiCycle Phase 2 constitute dense supervision that would otherwise be prohibitively costly to obtain by manual volumetric annotation, enabling scratch training of a DSM.

### Domain-specialized models enable 3D segmentation across diverse epithelial contexts

Having established DSMs for adult zebrafish epidermis, we next evaluated their ability to reconstruct full tissue-scale volumes within the original training domain (see Methods). Large-scale segmentation produced dense instance predictions spanning thousands of cells within the broader epithelial context (**Fig. 4A**). Qualitative inspection showed that many segmented instances exhibited plausible cell morphologies and preserved visually recognizable features of stratified epithelial organization, including flattened and anisotropic superficial keratinocytes and polygonal basal cells **(Fig. 4A1’-A3’; Video S1)**. These observations indicate that models trained on localized HITL-curated patches can be applied to larger volumes within the same biological context to generate tissue-scale instance predictions without additional supervision.

**Figure 4.**
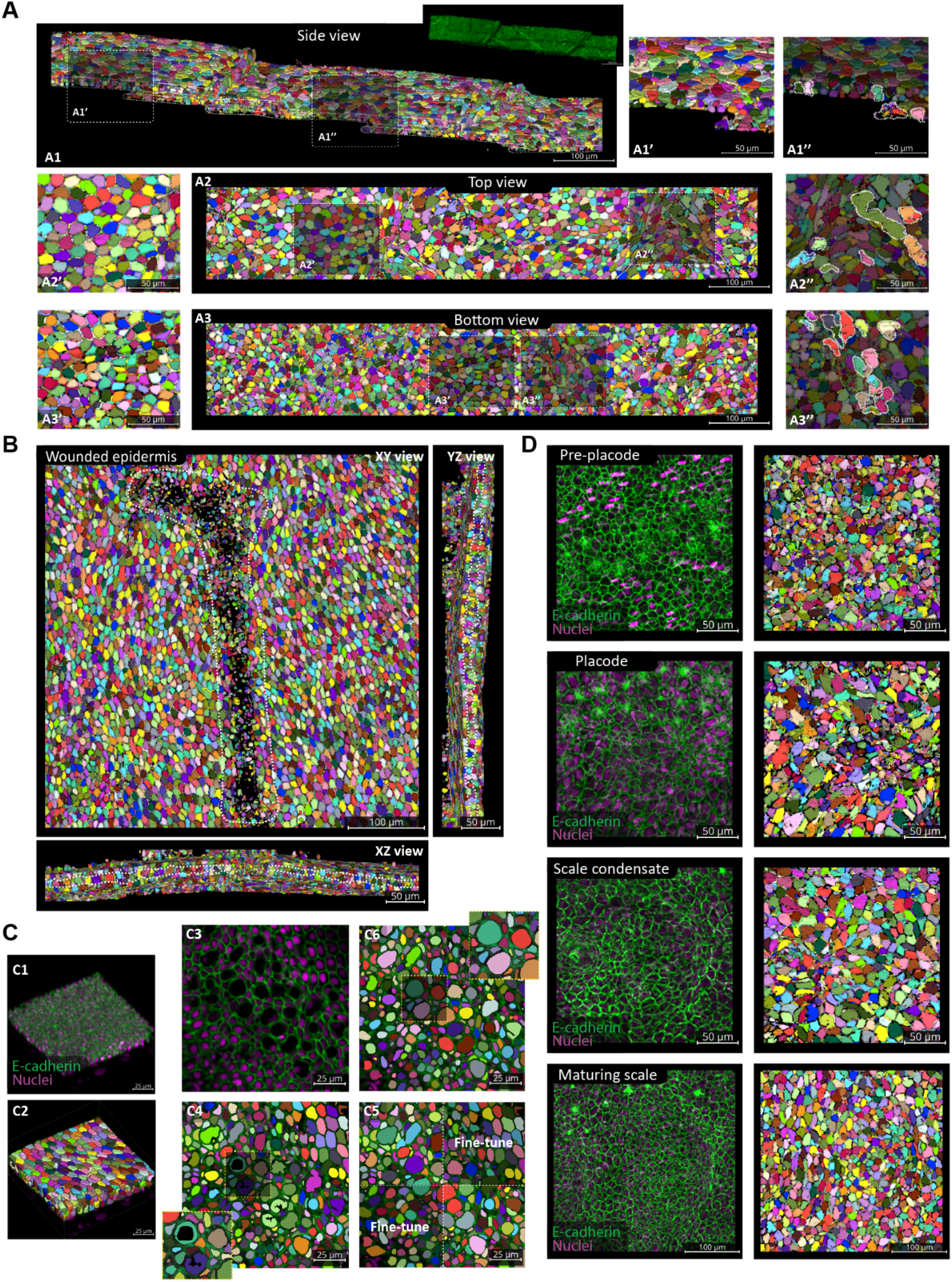
Domain-specialized model enables large-scale, cross-context epithelial segmentation. **(A)** Tissue-scale 3D reconstruction of adult zebrafish stratified epidermis showing large-area instance segmentation. Insets highlight representative local regions from different tissue depths and views (A1’,”: side views, A2’,”: top-down views and A3’,”: bottom views). Insets show representative regions of well-reconstructed cells (A1’-A3’) and poorly reconstructed cells (A1”-A3”). Outlines indicate areas of poor reconstruction and incoherent segmentations. **(B)** Segmentation of wounded epidermis. Lateral view shows a top-down view of wounded (inside outline) and unwounded regions. Axial views show stratified layers of the epidermis (outlines). **(C)** Local model refinement in a representative region of interest. (C1) Fluorescence volume and (C2) corresponding 3D instance reconstructions. (C3-C4) Representative mid-slice showing fluorescence and initial instance segmentation, respectively; inset in C4 highlights a region with poor reconstruction. (C5) Four local 3D sub-stacks selected for additional annotation and model fine-tuning; representative 2D slices are shown for visualization. Lateral and axial models were fine-tuned using corresponding XY and XZ/YZ plane samples from these sub-stacks. (C6) Segmentation of the whole region following fine-tuning, with inset showing the same area highlighted in C4 and improved resolution of previously poorly reconstructed cells. **(D)** Application of trained models to developmental stages of scale morphogenesis, including pre-placode, placode, scale condensate and maturing scale stages. Confocal images of the *TgBAC(cdh1:cdh1-sfGFP);Tg(actb2:h2b-2x-mScarlet)* transgenic line are shown alongside predicted instance segmentations. These examples illustrate model behavior across morphologically distinct epithelial states and highlight typical segmentation quality and failure modes outside the primary training stacks.

Because full-volume ground truth was not available at this scale, our evaluation was primarily qualitative. We note that segmentation quality was not uniform across the reconstructed volume. Regions with morphology and membrane signal characteristics like the training patches generally showed cleaner instance separation, whereas other regions exhibited localized errors, including undersegmentation, merged neighbor cells and boundary irregularities (**Fig. 4A1’’-A3’’; Video S1**). Segmentation artifacts were more frequent in deeper regions where membrane signal intensity diminished with imaging depth **(Fig. 4A1”,A3”; Video S1)**. This suggests that merges are mostly caused by weak image signal, not because the model fails outside the original training context.

We next tested whether the DSM could be applied to related epithelial contexts beyond the specific regions used for training. We applied the DSM to additional zebrafish epidermal datasets spanning injury and developmental contexts, which differed in epithelial thickness, curvature, cell morphology, packing density and membrane signal consistency. In tissues retaining broadly similar multilayered stratified organization, such as in the adult wounded epidermis **(Fig. 4B; Video S2)**, reconstructions frequently produced coherent instance masks under qualitative inspection, although localized segmentation errors remained present **(Figs. 4B; Video S2)**. In this context, we observed that certain enriched cell morphologies that were absent from the original training patches were associated with repeated undersegmentation or lower-quality boundaries **(Fig. 4C3, C4)**. To test whether such context-specific failure modes could be addressed through limited, targeted curation, we identified a representative subregion within the adult wound dataset exhibiting recurrent segmentation errors associated with underrepresented cell morphologies and reconstructed this region accordingly **(Fig. 4C1, C2)**. Following this, we refined the resulting reconstruction to correct for errors and performed an additional fine-tuning step using annotations from only a subset of the region, while withholding the remaining area **(Fig. 4C5)**. When we used the fine-tuned model to reconstruct the full tile, instance separation and boundary quality improved not only within the curated subregion but also in adjacent regions containing similar morphologies **(Fig. 4C6; Video S3)**. These results demonstrate that localized annotation within EpiCycle can selectively correct DSM failure modes without requiring extensive retraining.

We observed a related but more pronounced pattern in earlier developmental datasets, where segmentation errors and quality were strongly signal-dependent and exhibited systematic structure **(Fig. 4D)**. While basal cells often retained relatively strong boundary signals, other layers showed heterogeneous and locally discontinuous membrane labeling, leading to reduced instance resolution at cell-cell interfaces. Under these conditions, separation failures were enriched along the axial direction, where membrane boundaries between vertically adjacent cells were frequently incomplete, resulting primarily in undersegmentation and occasional multi-cell merging. As development progressed toward maturing scale stages, we observed that lateral cell boundaries became more consistently resolved and instance separation correspondingly improved **(Fig. 4D; Video S4)**. These stage-dependent trends are consistent with variation in membrane contrast and imaging quality across developmental timepoints, as well as differences in cell morphology and packing geometry relative to the adult training domain. Overall, these results indicate that DSMs can generate usable instance segmentation in related epithelial contexts without immediate retraining, but performance decreases as tissue morphology and imaging characteristics diverge from the original training data.

### Adapting EpiCycle models to reconstruct complex mammalian epithelia

To demonstrate EpiCycle’s generalizability to other complex stratified tissues, we applied the full workflow to two mammalian embryonic organs: the murine salivary gland and the human epidermis, both of which undergo extensive 3D morphogenetic changes during embryogenesis leading to complex tissue morphologies (Holbrook, 1979; Knosp et al., 2012).

First, we applied EpiCycle to an explant of embryonic murine salivary gland, an organ whose highly branched epithelial morphology diverges from the planar tissues our DSM was trained on. Within selected regions of the gland, we leveraged the full extent of the EpiCycle workflow to train separate lateral and axial model pairs from scratch to segment cells and nuclei, respectively (see Methods). The resulting final cell and nuclei model pairs were then applied across the whole tissue, yielding 82,063 cells and 79,582 nuclei, with 94.4% of cells (77,490) reconciled to at least one nucleus recapitulating the branched morphology of the gland with cells and nuclei that are, on visual inspection, qualitatively consistent with expected epithelial architecture across densely packed regions (**Fig. 5A,B; Video S5**).

**Figure 5.**
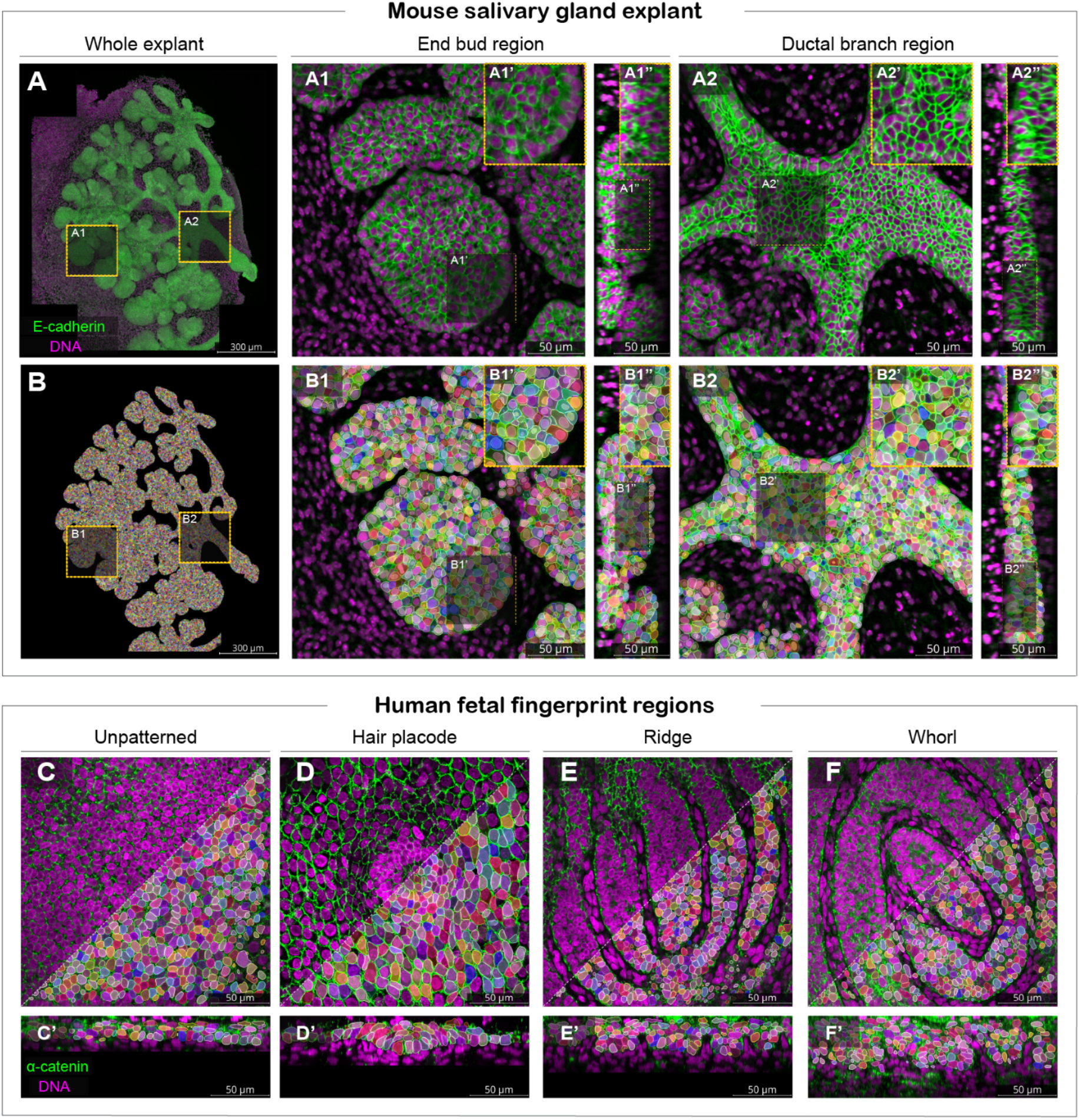
EpiCycle reconstructions of complex mammalian epithelia. **(A)** Raw confocal image of whole embryonic mouse salivary gland explant, with regions A1 and A2 indicating representative end bud and ductal branch regions, respectively. Enlarged midslice view of respective regions showing local epithelial architecture, with corresponding side views illustrating tissue organization throughout the z-axis. **(B)** Corresponding EpiCycle reconstruction of the salivary gland with enlarged mid slice view of end bud (B1) and ductal branch (B2) regions and respective corresponding orthogonal views illustrating cell and nuclei reconstruction across densely packed and highly irregular epithelial regions. **(C–F)** EpiCycle reconstruction of representative human fetal skin showing unpatterned epidermis (C), a hair placode (D), a fingerprint ridge region (E) and a fingerprint whorl region (F). En face views show mid-slice α-catenin and nuclei (DAPI) fluorescence together with corresponding individual cell instance reconstructions. **(C’-F’)** Corresponding orthogonal views illustrate reconstruction through the epithelial depth.

Second, we selected 13 confocal stacks of fixed human fetal digit and back skin spanning 12-14 weeks estimated gestational age (EGA), representing a range of epithelial architectures present during skin organogenesis (Glover et al., 2023) (**Fig. 5C-F; Table S1**). Notably, numerous microscopy parameters used during acquisition of this dataset differ from the zebrafish dataset. Starting from our zebrafish DSM, a single densely annotated 3D crop followed by [5/5] rounds of HITL fine-tuning across 29 crops spanning all 13 stacks yielded human-adapted lateral and axial model pairs, which reconstructed 3D cell masks across all specimens (see Methods). After quality control, the final reconstructions contained 17,436 cells across all reconstructed stacks for downstream analyses (**Fig. 5C-F; Video S6, representative samples**).

In sum, these applications illustrate EpiCycle’s flexibility in reconstructing complex epithelial architectures beyond zebrafish skin. By supporting both *de novo* model training and HITL-guided adaptation of existing DSMs, EpiCycle produces visually coherent cell (and nuclear) reconstructions across tissues that substantially differ in morphology, developmental origin, sample preparation and imaging conditions, demonstrating its broad applicability in datasets lacking prior ground truth annotations.

### EpiCycle reconstructions enable population-scale 3D morphometry in complex human epithelial tissue

As proof-of-principle of EpiCycle’s ability to generate biologically interpretable morphological data, we chose to focus on the human fetal skin dataset due to its diverse cellular and tissue architectures. From each retained cell instance, we extracted 31 morphological features spanning size, elongation, orientation, topology and surface asymmetry (**Fig. 6A; Table S3**). Applying principal component analysis (PCA) to these features identified interpretable axes of variation with the first three principal components (PC): PC1 (∼39% variance) represented cell scale, PC2 (∼16% variance) represented columnar height or thickness along the *z*-axis, and PC3 (∼13% variance) represented apical-basal cross-section asymmetry (**Fig. 6B,C**). The first 10 PCs accounted for ∼96% of the total variation and defined a 10-dimensional (10D) morphospace used for all downstream analyses. Hierarchical clustering of the PC loading matrix revealed that each morphometric axis draws on a distinct group of shape descriptors rather than redundant measures of a single property (**Fig. 6B**). To visualize the global organization of the morphospace, we projected the 10D morphometric coordinates onto two dimensions using UMAP and found a continuous morphological landscape rather than discrete islands (**Fig. 6C,D**). To delineate shape clusters within this continuum, we applied unsupervised leiden clustering and identified 16 clusters, each of which had characteristic shapes and showed enrichment for specific anatomical sites and donors (**Fig. 6D,E**).

**Figure 6.**
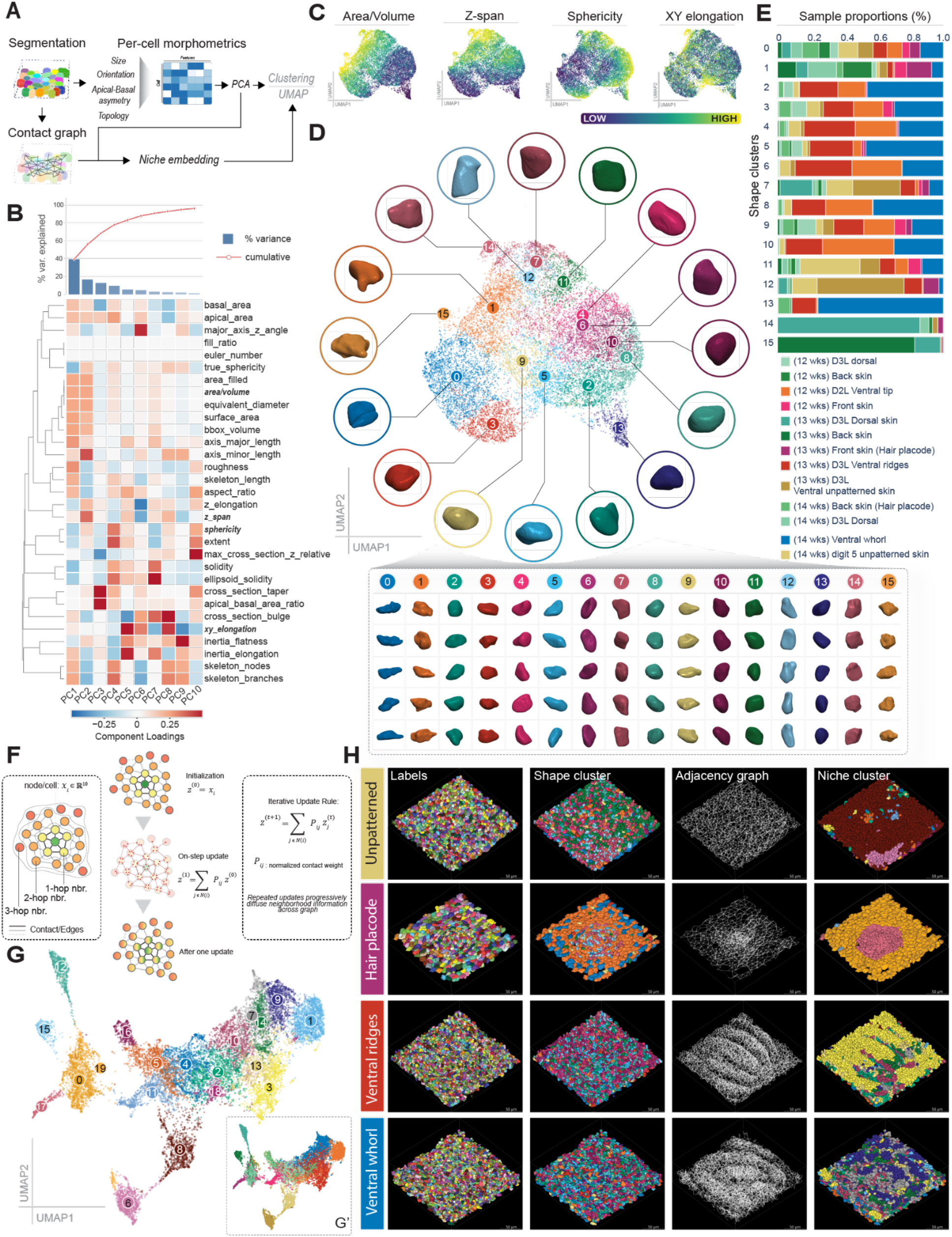
EpiCycle 3D reconstructions enable recovery of coherent cell morphologies and spatial tissue niches in human fetal skin. **(A)** Overview of the morphometric and spatial analysis workflow. From reconstructed 3D cell segmentations, per-cell morphological descriptors spanning size, orientation, apical-basal asymmetry and topology were assembled into a *Cell-by-Feature* matrix and reduced by principal component analysis (PCA, the Morphospace Embedding). In parallel, cell-cell contacts were used to construct a contact graph. Cell PC representations were subsequently combined with the contact graph to generate a neighborhood-level representation for each cell (termed as Niche Embeddings). Shape and niche representations were visualized by UMAP, and clusters were derived using the leiden clustering algorithm. **(B)** PCA of per-cell morphological descriptors. Top, variance explained by each component and cumulative variance across the first 10 PCs. Bottom, hierarchically clustered loading matrix illustrating contributions of individual features to each PC. **(C)** Representative morphometric features associated with each of the first four PCs projected onto the 2D UMAP of the 10D morphospace. Cells are colored by rank normalized feature values for visualization. **(D)** Unsupervised leiden clustering of the 10D morphospace. Representative 3D cell geometries are shown around the UMAP. Additional representative cells from each shape cluster displayed below (sizes are not drawn to scale). **(E)** Sample compositions of the 16 shape clusters across human fetal skin samples. Bars indicate the relative proportions of cells assigned to each cluster within each sample, highlighting differences in morphological composition across anatomical regions and donors. **(F)** Graph anatomy and construction of the niche embedding from cell morphology and physical tissue adjacency (left to right). Each cell is initialized as their computed 10D morphospace representation and iteratively blended with their neighbors as a weighted sum of their neighbor’s 10D representation for *N*-iterations. Each successive iteration allows incorporation of information from increasingly distant graph neighborhoods, with the final feature matrix (not shown) consisting of a new 10D representation. See methods for detailed graph construction and diffusion parameters. **(G)** UMAP plot of the resulting niche embedding, colored by leiden niche-cluster identity. Unsupervised clustering on niche embedding representation identified 20 clusters representing potential local combinations of cell morphology and neighborhood context. (G’) inset is the same UMAP representation colored by sample labels used in (E). **(H)** Mapping of shape and niche organization onto representative reconstructed human fetal skin regions. Columns show reconstructed, shape-cluster assignments, the physical cell-contact graphs and niche cluster assignments. Niche clusters form spatially coherent domains within the reconstructed tissues despite absence of positional or molecular information during clustering.

To ask how cell morphology is spatially organized within the tissue, we computed a niche embedding by diffusing each cell’s PC coordinates over its physical cell-contact graph (see Methods) (**Fig. 6F**), yielding a representation of the morphological profile of each cell’s immediate neighborhood. Dimensionality reduction and unsupervised leiden clustering within this space identified 20 niche clusters (**Fig. 6G**), which corresponded to visually interpretable spatial domains within the reconstructed tissue (**Fig. 6H; Video S6**). The resulting niche clusters were qualitatively robust across a range of niche diffusion and clustering parameters (see Methods) (**Video S7**). In several instances, niche clusters corresponded with visually recognizable tissue architectures, such as hair follicle placodes or fingerprint patterns, despite the absence of any molecular or positional priors during clustering (**Fig. 6H**).

We previously showed that basal cells of hair-bearing back skin and volar ventral digit skin have highly similar gene expression programs at 14 weeks EGA, with only a limited number of genes specific to each structure emerging at late stages of differentiation (Glover et al., 2023). Cell shape in that study was assessed in 2D sections, so we examined how 3D cell shape varied across anatomical sites. We found that cells from the same tissue occupied similar regions of the morphospace and formed a gradual transition from volar to hair-bearing skin (dorsal digit and front/back trunk) (**Fig. 7A**). Volar cells were generally smaller and more spherical (**Figs. 6C, 7A,C**), in line with the denser volar basal layer reported previously (Glover et al., 2023). Because the niche embedding averages morphology over each sample’s contact graph, some grouping by sample was expected. Indeed, the niche embedding grouped cells largely by sample with qualitatively modest overlap, and each niche cluster contained cells from several shape clusters (**Figs. 6G, 7B,D,E**). These observations suggest that tissue architecture is organized not only by which shapes are present, but also by how they are arranged.

**Figure 7.**
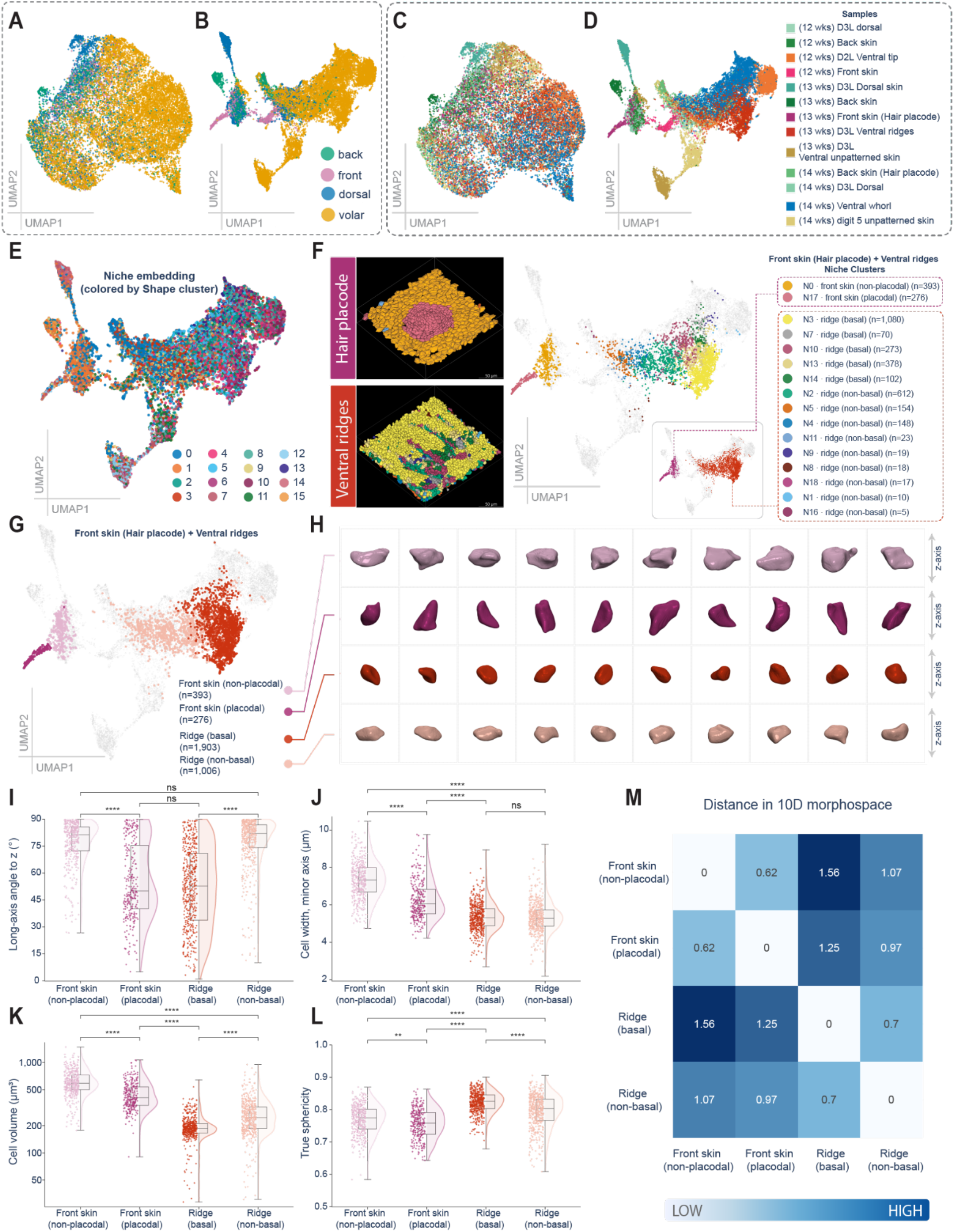
Anatomical organization of the cell morphospace and niche embedding, and comparison of hair placode and fingerprint ridge cell shape. (A–D) UMAP projections of the 10-dimensional shape morphospace (A, C) and of the niche embedding (B, D) for all 17,436 cells from 13 stacks, colored by anatomical site (A, B; back trunk, front trunk, dorsal digit, volar digit) or by sample (C, D; key at right, donor age in weeks EGA). **(E)** Niche-embedding UMAP colored by shape cluster (leiden, resolution 1.0, 16 clusters; Fig. 6D); most niche clusters draw cells from several shape clusters. **(F)** Cells used for the hair placode versus ridge comparison, from one donor at 13 weeks EGA. Left, 3D reconstructions of the front-skin stack containing a hair placode sample (top) and the ventral digit stack containing fingerprint ridges (bottom), colored by niche clusters (Fig. 6H). Right, niche-embedding UMAP with the niche clusters of the two stacks colored and all other cells in grey; inset, colored by the two samples. Hair placode cells are niche cluster N17 and ridge basal cells are N3, N7, N10, N13 and N14. **(G)** The same UMAP with the four comparison groups: front-skin epidermis, hair placode, ridge basal cells and the remaining cells of the ridge sample. **(H)** Representative cells of the four groups, rendered from the reconstructed 3D meshes. For each group, the ten cells closest to the group median in the 10-dimensional morphospace are shown, left to right by increasing distance from the median. All cells are drawn at a common scale, so sizes are comparable across rows; row colors as in G. **(I–L**) Distributions of four shape features per group: the angle between the cell’s principal (major/long) axis and the stack *z*-axis (degrees; 0°, parallel to the apical–basal axis; 90°, in the plane of the epithelium; I), cell width (minor axis length, µm; J), cell size (volume in µm³; K) and true sphericity (L). Points show individual cells (up to 500 per group, chosen at random). Boxes show the median and interquartile range (IQR), whiskers extend to 1.5 × IQR, and the half-violins show the kernel density. The box and violin plots were computed using all cells. Group sizes are given below each group. Groups were compared using Welch’s ANOVA with Games–Howell post hoc test on all cells (**, *P* < 0.01; ****, *P* < 0.0001; ns, not significant). **(M**) Separation of the four groups in the 10-dimensional morphospace: Euclidean distance between group medians divided by the mean within-group spread of the two groups, where a group’s spread is the median distance of its cells from the group median. Values above 1 indicate groups further apart than they are wide.

We next asked whether these tissue-level differences extended to individual epidermal structures. To eliminate potential confounds due to differences between individuals, we compared a hair placode (front trunk) and fingerprint ridges (ventral digit) at 13 weeks EGA from a single donor. Both structures corresponded to niche domains: the hair placode to cluster N17, and the ridge basal layer to clusters N3, N7, N10, N13 and N14 (**Figs. 6H, 7F,G**). We compared each structure with the remaining epidermis of its own sample and found that cells in both structures were oriented more along the apical–basal axis (i.e., *z*-axis) than their surroundings (**Fig. 7H,I**), but only placode cells were also narrower than their surroundings (**Fig. 7H,J**). Placode cells were larger than ridge cells, a difference also seen between the surrounding epidermis of the two samples (**Fig. 7K**), and ridge cells were rounder than placode cells (**Fig. 7L**). In the morphospace, the two structures occupied partly distinct regions (**Fig. 7M**). Thus, cells of the hair placode and fingerprint ridges differ in overall 3D shape, a distinction that EpiCycle makes measurable at single-cell resolution across intact tissue.

Together, these results demonstrate that EpiCycle’s bootstrap-derived 3D reconstructions alone retain sufficient geometric fidelity to enable unsupervised morphological analysis, recovering both meaningful morphospace and anatomically meaningful tissue domains without molecular information or prior curated labels.

## DISCUSSION

Accurate and scalable volumetric segmentation of cells in complex tissues remains a central challenge in biological image analyses, driven in large part by the scarcity of dense 3D ground truth and the substantial morphological heterogeneity observed across tissues and imaging conditions. In this work, we introduce EpiCycle to address this bottleneck by combining sparse 2D annotation, reconstruction-derived dense supervision and targeted model retraining. A key observation underlying EpiCycle is that dense 3D ground truth is not strictly necessary to achieve accurate volumetric reconstruction. Instead, we show that a modest amount of curated 2D supervision, introduced incrementally through HITL training, is sufficient to bootstrap reliable lateral and axial segmentation models. We show that EpiCycle enables the development of DSMs that scale from local regions of interest to whole-tissue volumes within a ground truth-limited setting across diverse epithelia. Our approach thus opens new avenues for accurate descriptions of complex tissue geometries, which has implications for understanding stratified epithelial development, regeneration and disease.

Recent methods such as u-Segment3D demonstrate effective aggregation of 2D segmentations into volumetric reconstructions (Zhou et al., 2025) but do not address how robust 2D models can be trained and refined in settings where dense 3D ground truth is currently unavailable or where models result in poor reconstructions. EpiCycle complements these approaches by explicitly closing the loop between sparse human annotation, reconstruction and model retraining/fine-tuning, thereby repurposing reconstruction as a source of dense supervision rather than solely an inference step. EpiCycle is instantiated using Cellpose-based 2D segmentation models and a modified version of direct aggregation method of u-Segment3D that allows support for user-trained Cellpose models and HITL refinement. While u-Segment3D supports multiple aggregation strategies, including indirect mask-based methods, the current EpiCycle workflow focuses on flow and probability-based direct aggregation due to its ability to enforce geometric consistency across slices. Conceptually, EpiCycle links Cellpose and u-Segment3D by introducing an explicit learning and refinement loop.

### Development and refinement of a domain specialized model

As a proof-of-principle to benchmark EpiCycle, we focused on the vertebrate skin epidermis, a stratified and highly ordered epithelium comprised of diverse cell geometries (Kubo et al., 2009; Mackenzie, 1975; Moll et al., 2005; Snell, 1967; Yokouchi et al., 2016; Zampetti et al., 1989). Keratinocytes undergo dynamic cell shape changes during differentiation and wound healing (Gault et al., 2014; Kennard and Theriot, 2020; Leybova et al., 2024; Martin and Lewis, 1992; Richardson et al., 2016; Sarate et al., 2024; Snell, 1967; Yokouchi et al., 2016). Keratinocyte geometry is tightly regulated during epidermal morphogenesis and homeostasis (Luxenburg and Zaidel-Bar, 2019; Sahu et al., 2025), with disruptions to keratinocyte shapes leading to impaired barrier function, hyperproliferation and inflammation (Koegel et al., 2009; Soffer et al., 2026; van der Kammen et al., 2017). Although the superficial location of skin makes it an attractive organ system for imaging, most existing methods for skin segmentation are either not based on fluorescent confocal imaging or not aimed at providing cellular-level resolution (Cai et al., 2020; Decencière et al., 2013; Lin et al., 2021; Prinke et al., 2021). We show that EpiCycle can robustly reconstruct keratinocyte geometries across organisms and imaging conditions.

We found that improvements in 2D segmentation quality translated directly into improved 3D reconstruction, thereby acting as an effective amplifier of sparse supervision **(Figs. 2,3)**. Notably, axial models saturated more rapidly than lateral models, likely reflecting both the larger number of axial training slices and relative ease of annotating axial cross-sections. These results provide a practical response to a long-standing challenge in volumetric segmentation: while *de novo* dense 3D annotation is costly and often infeasible, sparse 2D annotation is tractable and interpretable. EpiCycle leverages this asymmetry by explicitly structuring segmentation as a progressive learning process rather than a one-shot prediction task by allowing a model assisted labeling framework.

Our benchmarking results further demonstrate that variability in segmentation performance across stacks is primarily driven by morphological diversity rather than model instability or overfitting. HITL-trained bootstrap models achieve high median 3D AP@50 but exhibit substantial variability across sample tissues, likely reflecting exposure to rare or challenging morphologies. Instead, generating training data on fully reconstructed 3D volumes and subsequent training reduced this variability and improved worst-case performance, indicating that dense morphological coverage, rather than increased model capacity, is the dominant factor governing robustness. Importantly, patch-specific trajectories in median 3D AP@50 illustrated that tissues exhibiting initially poor performance often improved disproportionately once included in training, suggesting that many failure modes are learnable rather than intrinsic. Notably, a classical approach in the field is to fine-tune existing pretrained models within a given domain. However, as illustrated, fine-tuned Cellpose models failed to match models trained from scratch (**Figs. 3E, scratch; 3G, Cyto3 FT; S3**). We speculate that this likely reflects a difficulty in modifying the geometric priors embedded in Cellpose’s learned representation, whereas scratch training allows the model to learn priors and representations more suitable to our dataset. In this context, domain specialization should not be viewed as a limitation of the workflow but rather a form of convergence toward a stable solution within a user-defined biological and imaging regime.

Application of EpiCycle-trained models to full tissue volumes and related epithelial context showed both strong generalization and structured failure modes **(Fig. 4)**. DSMs generalized well to adult tissues with similar stratified organization and imaging appearance, producing coherent reconstructions without needing exhaustive annotation or training. By contrast, morphologically distinct contexts exhibited systematic errors, like under-segmentation of unusually large cells and failures in regions with markedly different illumination, likely reflecting underlying biological and imaging differences. Importantly, these failure modes were interpretable and morphology-linked rather than random, enabling targeted refinement. Within the EpiCycle workflow, such errors naturally motivate additional annotation cycles and selective fine-tuning, followed by reconstruction, allowing dense 3D supervision to be generated specifically for underrepresented morphologies. This targeted curation strategy reflects how segmentation models are refined in practice and contrasts with approaches that require training on densely annotated 3D datasets. Thus, EpiCycle enables the user to adaptively refine models in response to new failure modes.

### Intended usage and applications

EpiCycle is implemented as an open-source plugin for napari (Sofroniew et al., 2025). EpiCycle provides an integrated graphical, step-by-step interface that allows users to perform the complete workflow—from data selection through tissue-scale segmentation—within a single environment without requiring programming expertise (**Fig. S4**). A standard workflow begins by applying provided models to cropped regions of interest relevant to the user’s biological question. Based on qualitative inspection of the resulting 2D segmentations and 3D reconstructions, users may perform targeted HITL corrections, retrain or fine-tune models and adjust reconstruction parameters as needed. Once performance is satisfactory, the same models and parameters can be applied to larger tissue volumes with comparable imaging conditions, optionally leveraging shared computing resources for large-scale reconstructions.

The workflow emphasizes flexibility over prescription. Users may interleave sparse HITL rounds with dense consolidation, skip consolidation entirely or revisit model training as new data or failure modes arise. Dense fine-tuning is an optional consolidation step rather than a prerequisite for deployment—models can be used effectively after HITL rounds. Sparse HITL stages reduce correction burden over time, while dense consolidation provides incremental gains in reconstruction fidelity when required. Users are also not restricted to the provided models: the workflow supports training from scratch with user-defined annotations and segmentation ontologies, allowing task-specific definitions of cell boundaries, instances or cell types appropriate to the experimental system.

Beyond segmentation, EpiCycle integrates tools for downstream quality control and exploratory feature analysis. Per-cell features and quality control scores can be gated and explored interactively, with linked 3D inspection, preliminary dimensionality reduction and lasso-based selection. EpiCycle exports feature tables and adjacency graphs in standard formats, enabling downstream analyses (e.g., population-level analysis, joint morphospace construction or spatial niche embedding) in external tools. Our analysis of human fetal skin illustrates this end-to-end usage: DSMs trained on zebrafish were transferred to fixed human tissue via targeted fine-tuning, applied at scale across multiple stacks, quality-controlled interactively and analyzed as a joint 3D morphospace, progressing from raw confocal volumes to population-level shape analysis within a unified workflow.

In this setting, our comparison of hair placodes and fingerprint ridges relied on cell shape and spatial niche information, revealing a shared reorientation toward the apical–basal axis and placode-specific narrowing. Ideally, shape and gene expression would be measured in the same tissue, making it possible to directly ask how a cell’s 3D shape and position relate to its overall molecular identity or cell state. For example, one could ask whether placode narrowing accompanies the gene programs that distinguish placodes from ridges (Glover et al., 2023). Fluorescent in situ methods already enable quantitative, single-cell measurements of gene expression in whole-mount tissue (Choi et al., 2018), while spatial omics approaches are increasingly being extended from 2D sections to thick tissues (Schott et al., 2024; Sui et al., 2025; Wang et al., 2021b). Integrating these measurements with EpiCycle’s 3D segmentations could map transcripts to individual cells and relate molecular identity directly to cell shape and local tissue context.

The general scarcity in 3D ground truth presents a challenge to many segmentation applications, which has motivated recent novel 3D reconstruction strategies. For example, an emerging approach uses synthetic data in conjunction with sparse 2D mask labels to efficiently train native 3D models (Thiyagarajan et al., 2024), while u-Segment3D demonstrates that complementary 2D prediction can be resolved into accurate volumetric instances (Zhou et al., 2025). Complementary to these approaches, reconstructed volumes generated by EpiCycle could serve as pseudo-ground truth for learnable native 3D models. Lastly, we note that although we chose to focus on segmentation of stratified epithelium, EpiCycle could be applied to any segmentation objective in which expert curation improves model predictions.

### Scope and limitations

EpiCycle is designed as a domain-specialized workflow for 3D segmentation of stratified epithelial tissues. The following limitations should be acknowledged. First, as with all data-driven segmentation approaches, model performance depends on the representativeness of the annotated training data. Early HITL rounds that focus on spatially or morphologically homogeneous regions may bias the learned representations and delay exposure to rare configurations that ultimately affect the final 3D reconstruction. EpiCycle mitigates this through failure-driven iteration, in which systematic errors encountered post-3D reconstruction naturally prompt targeted annotation-curation and subsequent model retraining or fine-tuning cycles. Second, EpiCycle intentionally produces domain-specific models that perform robustly within a given biological, imaging and acquisition context, but does not guarantee zero-shot transfer to distant tissues or modalities without retraining or fine-tuning. Third, the workflow relies on high-quality 3D reconstruction to generate dense supervision. Thus, suboptimal reconstruction parameters or imperfect curations may propagate downstream errors into later rounds of training. Fourth, while EpiCycle substantially reduces the prohibitive labor associated with *de novo* dense 3D annotations, expert judgement is vital for identifying systematic failure modes, adjudicating ambiguous cases and defining task-relevant segmentation objectives.

## MATERIALS AND METHODS

### 1. Zebrafish husbandry and imaging

#### 1.1 Zebrafish lines and care

Zebrafish were housed at 26-27℃ on a 14/10 light cycle. The following transgenic lines were used: *TgBAC(cdh1:cdh1-sfGFP,cryaa:Cerulean)^sk95Tg^*(Colak-Champollion et al., 2019) and *Tg(actb2:h2b-2x-mScarlet)^hm63Tg^*(Peterman et al., 2025). Animals of either sex were used. All zebrafish experiments were approved by the Institutional Animal Care and Use Committee at the University of Washington (Protocol #4439-01).

#### 1.2 Confocal imaging

Live imaging of juvenile zebrafish was performed by anesthetizing animals in 0.006-0.012% buffered MS-222 diluted in system water. After 3 to 5 minutes, fish were moved to an imaging chamber, stabilized by covering with 1% agarose, and then covered in MS-222 diluted in system water. Images were captured using an A1R MP+ confocal scanhead mounted on a Ni-E upright microscope (Nikon). A 16×/0.8 water dipping objective (Nikon) with a 6x digital zoom applied was used for *z*-stack acquisition with a *z*-step of 0.25 μm and images were post-processed using the Denoise.ai function in NIS-Elements (Nikon). Acquisition settings for all datasets are described in **Table S1.**

For live imaging of adult zebrafish, an intubation-based protocol was used for image acquisition as in Xu et al. (2015). In brief, fish were anesthetized in 0.012% buffered MS-222 diluted in system water for 8-10 min until gill movement became very slow. Fish were transferred to a custom imaging chamber and embedded with 1% agarose, as described above. Continuous delivery of 0.08% buffered MS-222 into the mouth of the fish was achieved using polyethylene tubing (Becton Dickinson 427421 or 427400) and a peristaltic pump with a flow rate of 2-3 ml per min. Tiled z-stacks of the adult scale epidermis were acquired using a 16x/0.8 water-dipping objective (Nikon) with a 6x digital zoom using a *z*-step of 0.2 μm. To revive fish after image acquisition, the peristaltic pump line was transferred to a bottle containing system water and fish were closely monitored for recovery.

For imaging of the wounded epidermis, zebrafish were anesthetized and a scale was removed from the flank of the fish, placed on a 6mm plastic dish, and covered with L-15 media. The epidermis was mechanically injured using a pair of forceps, and tiled z-stacks were acquired using a 25×/1.1 objective (Nikon) at 4x digital zoom using a *z*-step of 0.2 μm and images were post-processed using the Denoise.ai function in NIS-Elements (Nikon).

### 2. Mammalian samples

#### 2.1 Human fetal skin

Human fetal tissue samples were obtained after elective medical termination of pregnancy from the Royal Infirmary of Edinburgh, UK with informed consent (approved by the Lothian Research Ethics Committee, ref. 08/S1101/1). Only morphologically normal samples were included and samples from both sexes were used. Gestational age was estimated by ultrasound scan before the procedure and confirmed by measuring foot length afterward and reported as weeks EGA. Human fetal skin samples were stained and imaged as previously described (Glover et al., 2023). Briefly, formalin fixed skin samples were washed in PBS containing Triton X-100 (PBTX, Sigma-Aldrich), blocked for 2 hours in 10% goat serum/PBTX at room temperature, before incubation with anti-α-catenin antibody (Santa Cruz Biotechnology #sc-9988) diluted 1:100 in blocking buffer for 24 hours at 4°C. Samples were then washed in PBTX before incubation with goat anti-mouse IgG Alexa Fluor 488 secondary antibody (1:500, Life Technologies # A11029) and Alexa Fluor 647 Phalloidin (1:200, Life Technologies #A22287) diluted in blocking buffer for 24 hours at 4°C. Samples were washed in PBTX, counterstained with DAPI/PBTX for one hour at room temperature and mounted in ProLong Gold. *z*-stacks of dimensions 1024 × 1024 were acquired on whole mount digit-stained skin samples using a *z*-step of 1 μm and an LSM 880 confocal microscope (Zeiss) equipped with a 40×/1.3 PL APO oil objective.

#### 2.2 Mouse salivary gland explant preparation and image acquisition

All mouse experiments were conducted under the approval of the ethics committee of the Institut Curie and the French Ministry of Research (reference #34364-202112151422480) in the Animal Facility of Institut Curie (facility license #C75–05–18). Salivary glands were dissected from mouse embryos at E14.5 days of development as previously described (Wang et al., 2021a). Briefly, embryos were collected from the uterus of pregnant females and stored on ice in 24-well plates. Collected salivary buds with surrounding mesenchyme were placed on a cell culture insert floating on culture medium within a 6-well plate. The embryonic culture medium consisted of DMEM/F-12 supplemented with 1x GlutaMax, 10 mM HEPES, 2% insulin-transferrin-selenium and 2% penicillin-streptomycin (**see Table S4 for catalog numbers**). Explant cultures were grown in a tissue culture incubator at 37°C with 5% CO_2_.

Following tissue isolation, whole mount immunostaining was performed on the explants at the end of the culture (48 hours). Tissues were fixed in 4% PFA for 15 minutes at room temperature (RT), permeabilized with 10% (w/w) Triton X-100 in PBS at 4°C overnight and blocked in 5% FBS, 1% BSA in PBS for 1 hour at RT. Samples were incubated with a rabbit anti-E-cadherin primary antibody (Cell Signaling Technology Cat# 3195, RRID:AB_2291471) at 1:200 for 48 hours at 4°C. After removal of the primary antibody solution and 3x washes in PBS, samples were stained overnight at 4°C with a goat anti-rabbit Cy5-conjugated secondary antibody (Thermo Fisher Scientific Cat# A10523, RRID:AB_2534032) at 1:300 and DAPI (Merck Millipore, CAS 28718-90-3) at 1:1000. After 3x washes in PBS, explants were mounted on a microscope slide using Aqua-Polymount.

Images were acquired using an LSM780 inverted laser scanning confocal microscope (Zeiss) equipped with a 40×/1.3 DICII PL APO oil objective. Tiled *z*-stacks were acquired using a *z*-step of 0.5 μm.

#### 2.3 Data preprocessing

All image preprocessing steps were performed in ImageJ/Fiji (Schindelin et al., 2012). Confocal stacks were 3D Gaussian blurred with default parameters (*σ_zxy_* = (2.0, 2.0, 2.0)). Subsequently, each stack was then rescaled to isotropic voxel sizes and background subtraction was performed with default rolling ball radius parameters (*r* = 50 *pixels*).

### 3. Ground truth generation and training data preparation

#### 3.1 Ground truth curation

To develop and benchmark EpiCycle, eleven densely annotated 512 × 512 × *Z* volumetric ground truth datasets from adult zebrafish trunk epidermis were prepared, eight of which were cropped from tissue-scale acquisition described in §6.1. For each stack, initial 3D instance predictions were generated using naively fine-tuned Cellpose models (trained in the Cellpose GUI following the developer-recommended workflow) (Stringer et al., 2021) applied to lateral and axial views and aggregated using the indirect reconstruction method of u-Segment3D (Zhou et al., 2025). All resulting 3D instance masks were subsequently curated and corrected in 3D using the EpiCycle multiview annotator feature within napari. Seven stacks were assigned as model-development data (patches 1-5, 7-8) while remaining four stacks (patches 6,9,10 and 14) were designated as test stacks and were not seen by any model during 2D bootstrap HITL training experiments illustrated in Figure 2. Models trained for dense supervision as illustrated in Figure 3 and Figure S3 used exclusively the development data or all eleven stacks accordingly.

#### 3.2 Isotropic resampling and orthoview decomposition

Each curated stack consisted of multichannel volume *I(z,c,x,y)* and a corresponding 3D instance label volume *M(z,x,y)*, where *z* indexes depth, *c* indexes fluorescence channels (membrane and nucleus), and (*x, y*) denote the lateral dimensions. Prior to slice extraction, all volumes were resampled to isotropic voxel spacing using ImageJ/Fiji.

To enable multiview learning of tissue morphology from 2D inputs, curated 3D volumes were decomposed into orthogonal 2D slice sets corresponding to lateral (*xy*) and axial (*xz, yz*) views. Lateral slices were obtained directly along the z-axis as *I(z,c,x,y)* with corresponding labels *M(z,x,y)*. For axial views, volumes were transposed so that slices were indexed along the appropriate dimensions, yielding slice stacks *I(y,c,z,x)* and *I(x,c,z,y)* for *xz* and *yz* views respectively. This decomposition preserves biologically meaningful morphological features of stratified epithelia: lateral slices capture epithelial tiling, boundary geometry, and in-plane cell curvature, whereas axial slices enable a trained model to learn apical-basal stratification, variation in cell height and width and cell packing across the epithelial depth.

#### 3.3 Slice filtering and mask refinement

The axial slice stack contains regions at the beginning and the end of each stack with incomplete tissue coverage where instance masks are sparse, absent or malformed due to edge effects during the 3D curation process. To avoid training on such negative slices and reduce background class imbalance, an edge buffer of 50 slices was excluded from both ends of each axial slice stack prior to training. Lateral slices were not buffered, so that the model also learns which layers contain no cells; empty lateral slices were retained in training.

Instance labels for each 2D slice were further refined prior to training. For each 2D slice, boundaries between touching cells were removed, connected components were relabeled, and instances smaller than 100 pixels were removed to eliminate debris and spurious fragments to ensure that spatially disconnected regions were treated as separate instances. Remaining labels were smoothed using morphological opening and closing with a disk structuring element (radius 1px) to standardize boundary geometry and improve consistency between slices.

#### 3.4 Training/validation/test splits and aggregation across stacks

For each curated 3D stack and for each view orientation, slice indices were partitioned into training, validation, and test splits using a spatially aware chunked splitting strategy. Specifically, slices were shuffled within chunks and allocated to splits in fixed proportions (40%/30%/30% and 80%/20%/0% training/validation/test for data conservative and data permissive training regimes respectively). This scheme preserves spatial contiguity within blocks while avoiding overly optimistic evaluation due to adjacently correlated slices that may leak across splits.

After per-stack splitting, slice groups were aggregated across stacks. For lateral training, only *xy* slices were pooled; for axial training, both *xz* and *yz* slices were pooled into a single dataset. Each slice was represented as a multichannel 2D image tensor (*c,x,y*) paired with its corresponding 2D instance mask *M(x,y)*.

### 4. EpiCycle model training

#### 4.1 Bootstrap HITL training

To evaluate the effectiveness of iterative HITL training for 2D cell segmentation, a simulated bootstrap training regime was implemented using seven model-development ground truth stacks (patches 1-5, 7-8). The four held-out stacks (patches 6,9,10,14) were not used during HITL training at any round.

At each training round, 15 lateral (*xy*) and 30 axial (15 *xz* + 15 *yz*) slices per stack were randomly sampled without replacement, yielding 105 lateral and 210 axial slices per round across the seven stacks. To avoid sampling uninformative regions at stack boundaries, the 50-slice axial edge buffer described in §3.3 was applied during sampling; no exclusion was applied to lateral slices. A minimum inter-slice distance of 5 slices was enforced between sampled slices within each stack to reduce spatial correlation. Sampled slices were accumulated across rounds, so the accumulated training set grew linearly from 105 lateral/210 axial slices at round 0 to a maximum of 700 lateral/1,400 axial slices once the available pool was exhausted (**rounds 6-7; Fig. S2**). Training data and hyperparameter configurations for each model in this study are listed in **Table S5**.

Prior to training, instance labels for each sampled 2D slice were refined as described in §3.3 (connected-component relabeling, removal of instances <100 px, morphological opening/closing with a disk structuring element of radius 1 px). Separate lateral and axial models were trained at each round. At round 0, both models were initialized from scratch (no pretrained weights). For rounds 1-9, each model was started from the previous round’s checkpoint and fine-tuned on the cumulatively expanded training dataset. All models were trained for 1,000 epochs using stochastic gradient descent with learning rate 1 × 10^−2^, weight decay 1 × 10^−5^, momentum 0.9 and batch size 32. Two input channels were used, corresponding to membrane and nuclear fluorescence signals. Images were normalized and rescaled internally by the Cellpose python package. Simulated HITL training was performed on an NVIDIA H200 GPU. Ten rounds of training yielded ten lateral and ten axial model checkpoints together with their corresponding cumulative training datasets for subsequent cross-round evaluation.

#### 4.2 Domain-specialized model training

Following the bootstrap HITL protocol, a pair of DSMs were trained suitable for application on novel data from the same tissue context. Starting from the round-4 HITL checkpoint (corresponding to early performance saturation; see §5.1), lateral and axial models were further fine-tuned on dense annotation aggregated across all eleven curated 3D ground truth stacks, using the slice decomposition and splits described in §3.2-3.4.

For lateral DSM, all *xy* slices from the training split were used; for the axial DSM, *xz* and *yz* training slices were pooled. Fine-tuning on the training data used two input channels (membrane and nucleus) and default Cellpose loss functions with stochastic gradient descent. Lateral and axial models were fine-tuned for 500 epochs, with batch size 16, learning rate 1 × 10^−2^, weight decay 1 × 10^−5^ and momentum 0.9 (**Table S5**). Validation splits were used for monitoring stable loss convergence and to detect overfitting but were not used for hyper parameter tuning or model selection, because 2D slice-level losses do not correspond directly to downstream 3D reconstruction performance. Final model selection was therefore performed exclusively at the 3D reconstruction level (see §5.2).

Benchmarking-scale training and large-scale inference were performed on NVIDIA A100 or H200 GPUs. Models used for demonstrating typical laboratory usage were trained on a single NVIDIA RTX 4000 GPU.

#### 4.3 Ceiling (fully supervised) model training

To estimate the performance ceiling achievable under dense volumetric supervision, a pair of lateral and axial models were trained from scratch (no pretrained weights) using the fully dense annotations from all eleven curated 3D ground truth stacks (or exclusively the seven development stacks). Training hyperparameters otherwise matched those described for DSM training in §4.2 (500 epochs lateral and axial, batch size 16, learning rate 1 × 10^−2^, weight decay 1 × 10^−5^, momentum 0.9, two input channels, default Cellpose loss functions and conservative or permissive training data regimes; **Table S5**). These fully supervised scratch models serve as an upper bound on the reconstruction accuracy achievable within this tissue imaging context.

#### 4.4 Cellpose cyto3 fine-tuning comparison

To assess whether a pretrained generalist Cellpose model could be adapted to our tissue context, pairs of lateral and axial models from Cellpose cyto3 checkpoint (Stringer et al., 2021) were fine-tuned on the same dense training data for DSM and ceiling training. Fine-tuning was performed at two learning rates: 1 × 10^−2^ and 1 × 10^−1^ (**Table S5**). All other training parameters matched those of DSM and ceiling model training (§4.2-4.3).

### 5. Model evaluation

#### 5.1 2D cross-round model evaluations

To characterize how 2D segmentation performance evolved across HITL rounds, a pairwise cross-round evaluation was performed in which each of the ten lateral (and ten axial) model checkpoints was evaluated on each of the ten cumulative training datasets, yielding a 10×10 evaluation matrix per view.

For each model-dataset combination, predictions were generated with the `CellposeModel.eval()` function from the Cellpose python package and the membrane channel as the primary channel to segment and the nuclei channel as the secondary complementary channel utilizing default segmentation parameters accordingly (normalize set to True, cell probability threshold set to 0.0, flow threshold set to 0.4, diameter set to None and rescale set to True (default Cellpose parameters). Inference was performed in batch mode on an NVIDIA H200 GPU.

Per-slice segmentation quality was assessed using average precision at 50% intersection-over-union (AP@50), computed via ‘cellpose.metrics.average_precision’. AP@50 was chosen as the primary metric because predicted 2D masks serve as intermediate inputs to 3D reconstruction via probability and flow-field integration, where approximate instance localization matters more than pixel-precise boundary identification.

For lateral evaluation, slices that produced a degenerate zero score with no overlap between prediction and ground truth (dice = 0 and AP@50 = 0; 4,046 of 50,050 slices, 8.1%) were excluded from the AP@50 summaries shown; these are dominated by empty edge slices at the top and bottom of the confocal z-stacks (no tissue) but also include slices where a model produced no output. Slices with partial overlap that failed the IoU-0.5 match or genuine detection failure (dice > 0, AP@50 = 0; *n* = 310) were retained. On the final (round-9) dataset this left *n* ∼ 634-659 lateral slices per model. The round-over-round improvement was robust both to this exclusion and to no filtering (median AP@50 shifted by <= 0.02). Axial slices always intersect tissue and are unfiltered (*n* = 1,400).

#### 5.2 3D reconstructions and benchmarking

Trained lateral and axial models were applied independently to 3D confocal volumes to produce per-slice 2D instance predictions across each available view using default Cellpose prediction parameters. Lateral *xy* predictions consisted of serial 2D segmentations *I*(*z*, *c*, *x*, *y*) → *M*(*z*, *x*, *y*); axial predictions consisted of *I*(*x*, *c*, *z*, *y*) → *M*(*x*, *z*, *y*) and *I*(*y*, *c*, *z*, *x*) → *M*(*y*, *z*, *x*) for *yz* and *xz* slices, respectively, producing per-slice probability and flow field predictions within each orthoview. Slice-level predictions from all three (*xy*, *xz*, *yz*) views were aggregated into 3D consensus instance labels using a modified version of the *direct* reconstruction method of u-Segment3D (Zhou et al., 2025), in which object probability maps and flow fields are integrated across 3 orthogonal views and volumes. Modifications to the u-Segment3D direct method implementation consisted of: 1) the ability to accept custom user-trained Cellpose model checkpoints as inputs; and 2) anisotropic pre-smoothing of raw probability maps (Cellpose’s per-pixel estimate of how likely a pixel is inside a cell) for each view’s segmentations to resolve possible noisy predictions for each independent slice–e.g., *σ_zxy_* = (3.0, 1.0,1.0) in which stronger smoothing is applied along the *z* axis for per slice *xy* segmentation results–prior to reconstruction. Pre-smoothing was used for the tissue-scale reconstructions (§6) but was not applied in the benchmark (**Table S6**).

Reconstructed 3D segmentation was evaluated against expert-curated 3D ground truth using 3D average precision at 50% IoU (3D AP@50), computed per stack and aggregated across datasets. 3D IoU was computed voxel-wise per instance pair using pre-defined metrics provided by the u-Segment3D package. This metric reflects the downstream biological task of volumetric cell instance segmentation and serves as the primary quantitative comparison between models. Across all comparisons, reconstruction parameters were held fixed across benchmarking experiments to isolate the effects of 2D model quality and downstream reconstruction performance (**Table S6**).

### 6. Large-scale and cross-context applications

#### 6.1 Tissue-scale reconstructions

Tissue-scale cell and nucleus reconstructions were produced by running the same pipeline described in §5.2 utilizing appropriately trained or fine-tuned lateral and axial model pairs. To facilitate this process, a tiling-based approach was used to partition the global sample data into manageable chunks. Specifically, the volume was divided into overlapping tiles of prespecified fixed dimensions and tile overlap (typically 1024 × 1024 × *Z* and 25% tile overlap along the imaging planes). To handle data of any dimension, the last tile was shifted inwards, which resulted in the last tile sharing larger overlap than initially prespecified. This approach avoided exposing the models to artificial boundaries, circumvented the need for artificial padding and ensured that the reassembled volume was exactly the size of the original input image. Within each tile, the tissue-containing depth-range was identified by the span between the first and last *z*-slice exceeding the mean brightness of 0.07 in a prespecified reference channel (e.g., GFP/membrane or DAPI/nuclei).

Each tile was then cropped in the *z*-dimension to the tissue-containing depth-range and 2D segmented with the appropriate lateral and axial model pair (*xy* slices for lateral models, *xz* and *yz* for axial models) for a given channel with the default Cellpose parameters noted in §5.1. The resulting corresponding probabilities and flow outputs were then used for consensus 3D reconstructions as described in §5.2. Afterwards, the resulting 3D instances were postprocessed by filling internal holes and discarding pieces below a minimum size and morphological operations (isotropic erosion, dilation or smoothing) and the original dimensions of the tile were recovered.

To recover the full global reconstruction, the per-tile 3D reconstructions were stitched into a single whole-tissue segmentation. Because neighboring tiles overlapped, the same cell was usually segmented in two adjacent tiles, or more at grid corners. These duplicates were identified by measuring how much each pair of overlapping cells coincided, using the intersection-over-union (a standard overlap score from 0 for no overlap to 1 for identical shapes), and treating two cells as the same when they overlapped by at least 0.25 and shared a minimum number of voxels. Each cell was linked to only its single best-overlapping counterpart in a neighboring tile, a conservative choice that errs towards keeping cells separate rather than merging them, and chains of linked overlaps (cell A to cell B, cell B to cell C) were combined into one cell. Overlaps were computed in blocks along the depth axis, so the whole volume was never held in memory at once. Each tile was then made responsible for the region up to the midpoint of its overlap with each neighboring tile, so that every voxel was written by exactly one tile. For potential downstream quality control, each final cell was labeled by how many tiles it spans (interior, spanning one seam, or spanning a corner) and by its weakest overlap score, giving a ranked list of the joins least likely to be correct.

Finally, the stitched segmentation was cleaned in three steps: 1) instances below a small voxel threshold (segmentation debris) were removed; 2) small clusters of instances detached from the main tissue mass were removed, identified on a coarsened occupancy map and discarding any disconnected components below a set fraction of the largest; and 3) each instance was surface-smoothed isotropically (*σ* = 1.5).

For the work presented here where cell and nuclei reconstruction was performed, cell and nucleus segmentations were matched as follows. Both results were produced from the same volumetric data on the same tiled grid and therefore aligned voxel for voxel. The reconciliation treated the cell segmentation as correct and adjusted the nuclei to match it. Each nucleus was assigned to a cell that contained the majority of its voxels. A nucleus that spanned the boundary between two cells was divided along that boundary, and each cell retained its own portion as a separate nucleus; the cell holding the larger portion retained the original label, and the remaining portions were relabeled accordingly. Portions smaller than 20% of the nucleus were discarded as misalignment between the two segmentations. Divided nuclei were then surface smoothed isotropically (*σ* = 1.5). Cells with no assigned nucleus were re-examined against the nuclear channel by percentile-based thresholding (p99 - p50 > 3 x (p50 - p16)) gated on whether in-cell contrast was sufficient to recover nuclear signal. The largest connected component above a threshold set midway between the cell median and its brightest voxels (p50 + p99)/2 was recovered as a nucleus if it was >= 10 voxels, <50% of the cell volume, and interior. Cells that still contained two or more nuclei were retained and recorded and were not divided.

All processing was performed on a high-performance computing cluster (HPC) with parallelization across multiple GPUs and CPU workers. Exact model pairs, tile dimensions, overlaps, thresholds and postprocessing configurations vary by dataset (**Table S6**). For zebrafish data presented in this work, all parameter configurations were kept identical and no post-stitching cleanup was performed, as this reconstruction was intended to demonstrate feasibility rather than to support quantitative morphometric analyses.

#### 6.2 Targeted fine-tuning on wound data

Wound data was processed in the same manner detailed in §6.1. Afterwards, to assess whether localized annotation could correct context-specific failure modes, a representative 1024 × 1024 × *Z* subregion exhibiting segmentation errors associated with underrepresented cell morphologies was identified within the adult wounded epidermis dataset. The initial DSM reconstruction of this subregion was manually corrected using EpiCycle’s multiview annotator. A subset of the corrected annotation was prepared to fine-tune the DSM using the methods described in §3.3 and §3.4 (80%/20%/0% training/validation/test splits after 3D→2D decomposition; **Table S5**), while the remaining annotated area was withheld as an internal evaluation region as illustrated in **Fig. 4C**. The fine-tuned model was then applied to the full tile, and instance separation and boundary quality were assessed qualitatively within (i) the curated subregion used for fine-tuning, (ii) the withheld annotated region, and (iii) adjacent regions of the full tile containing similar morphologies but no targeted annotation. Fine-tuning hyperparameters otherwise matched those of DSM training (§4.2).

#### 6.3 Human fetal skin transfer learning

To apply EpiCycle to human fetal epidermis, the adult zebrafish DSM was adapted using a two-step training approach. First, the zebrafish DSM was fine-tuned on a small 3D annotated 256 × 256 × *Z* cropped region of a single human fetal stack in which 2D training data was derived using the same methods detailed in §3.2-3.4 yielding (127/31/1) lateral (*xy*) and (204/51/1) axial (*xz* + *yz*) training/validation/test 2D ground truth slices, producing an intermediate human-adapted model pair. Secondly, these model pairs were then used to initialize HITL loop training over 29 additional 512 × 512 × *Z* crops sampled across all 13 human fetal stacks, thus ensuring that the models were exposed to the full range of acquisition conditions and epithelial architectures present in the dataset. At each round, model predictions were presented for correction in EpiCycle’s annotation interface; slices for which predictions were incorrect or unusable were re-annotated *de novo*. Corrected slices were accumulated cumulatively across rounds with checkpoint inheritance in the same manner described in §4.1. In sum, lateral and axial models were trained over 5 rounds, producing a final cumulative training set of 121 lateral and 53 axial ground truth slices. This two-step strategy reduced annotation burden of direct cross-species transfer by progressively exposing the model to larger and more diverse regions of the target domain in a way that was manageable for the user. The final model was then applied to all human fetal stacks (**Table S1**; model training in **Table S5**) to produce 3D cell instance reconstructions as described in §5.2 for downstream morphological analysis.

#### 6.4 Mouse salivary gland explant whole tissue reconstructions

To apply EpiCycle for volumetric reconstructions of mouse salivary explant dataset, cropped regions of interest (ROIs) (1024 × 1024 × *Z*) were preselected for 2D HITL training. The zebrafish DSM model was used to generate the initial lateral and axial 2D masks for training. Subsequent 2D HITL training resulted in 16 and 18 sparse GT-lateral and GT-axial slices, respectively. Subsequently, a smaller subset of 512 × 512 × *Z* patches were selected to reconstruct and curate within EpiCycle’s curation step (remove small objects, split merged instance, per-instance isotropic smoothing followed by morphological operations, including dilation). These curated patches were then used to prepare data and train final model pairs as described in §3.3 and 3.4, with the hyperparameters listed in **Table S5**. The resulting models were then used for full 3D reconstructions of whole salivary gland tissue using the same procedure detailed in §6.1 with tile size: 1024 × 1024 × *Z* and tile overlap: 25%, which resulted in 42 tile (7×6) grid partitions (**Table S6**). All Cellpose parameters used for predictions (cell probability threshold, flow threshold, diameter and rescale) were set to the default parameters.

For nuclei segmentation models, pre-existing Cellpose nuclei models were used to obtain initial mask predictions on hand selected slices within the same preselected ROIs. Predictions were manually curated for initial 2D predicted slices for initial 2D HITL on the same preselected ROIs used for cell segmentation resulting in 14 and 12 sparse GT-lateral and GT-axial slices, respectively. Subsequent final model training followed the similar procedure as cell segmentation model training.

Independent raw cell and nuclei reconstructions produced 90,862 cells and 87,411 nuclei 3D instances respectively. After removing debris and detached fragments (§6.1), 82,165 cells and 80,169 nuclei remained. Cell and nucleus instances were then matched (§6.1), resulting in a final reconciled set of 82,063 cells and 79,582 nuclei, of which 77,490 cells (94.4% assigned a nucleus) contained at least one nucleus and 4,573 (5.6%) contained none (**Fig. 5B**, reconstructions). Dividing nuclei that spanned a cell boundary restored a nucleus to 3,035 cells that would otherwise have been recorded as having none, and re-examination of nuclear staining recovered an additional 269 nuclei. Further 1,867 cells (2.3%) contained two or more nuclei and were retained. Lastly 4,163 nuclei fell outside every cell and were discarded.

### 7. Morphological analysis

#### 7.1 Shape feature extraction and cell-cell contact graphs

For each reconstructed 3D cell instance, EpiCycle extracts quantitative features in nine groups: core morphology, shape and orientation, surface, skeleton topology, cross-section profile, intensity, membrane and boundary intensity, cell–cell adjacency and probability consensus **(Table S3)**. Features are computed in a tiered fashion based on the availability of optional inputs: 19 morphology, shape, and orientation features are always computed from the segmentation masks alone; surface, adjacency, and topology features additionally require the label volume; intensity and membrane features require the source fluorescence image; and 12 probability-consensus features are extracted from the resulting u-Segment3D fused probability maps.

Core morphology and inertia-derived shape/orientation features were computed from the 3D label mask using `scikit-image.measure.regionprops_table` together with derived bounding-box and inertia-tensor statistics, yielding volume, equivalent diameter, extent, euler number, principal-axis lengths, elongation, flatness, and major-axis orientation descriptors. Surface descriptors (surface area, true sphericity, roughness, solidity) were computed from local 3D mesh or convex-hull reconstructions of each mask. Cross-section profile descriptors (apical and basal areas, apical-basal area ratio, maximum cross-section z-position, cross-section taper, and cross-section bulge) were computed from the area of each cell’s mask in every z-slice it spans. Intensity and membrane features summarize within-mask signal level and boundary-versus-interior contrast from user-selected image channels, while probability-consensus features quantify confidence, entropy, and boundary sharpness from the fused u-Segment3D probability volume (**Table S3**). Features that are numerically redundant with other measured features (e.g., compactness with extent) are computed but excluded from the morphospace feature set (§7.4).

A cell-cell contact graph for each stack was derived by graduated binary dilation of each cell mask: each cell’s mask was incrementally dilated up to a maximum radius (*r* = 5), and the first-contacted neighbor label together with the number of shared (overlapping) voxels was recorded. The resulting table, exported as `{stack_name}_adjacency.csv` with columns `[label_a, label_b, contact_voxels]`, provides an undirected, voxel-area weighted approximation of the physical surface contact between adjacent cells that served as the input to the niche embedding analysis (§7.6). All settings used for feature-extraction and generation of contact graph are listed in **Table S6.**

#### 7.2 Composite quality-control scoring

A per-cell composite quality-control (QC) score was computed as the mean absolute robust z-score across 15 curated features selected to minimize multicollinearity (**Table S3**). Prior to standardization, right-skewed unbounded features were log-transformed using a curate-plus-automatic scheme based on feature identity and empirical skewness (*x* ↦ *log*(*1* + *x*) where applicable). Each feature was then standardized using robust statistic (median and median absolute deviation, MAD), and the resulting z-scores were winsorized at ±10 to limit the influence of extreme outliers:

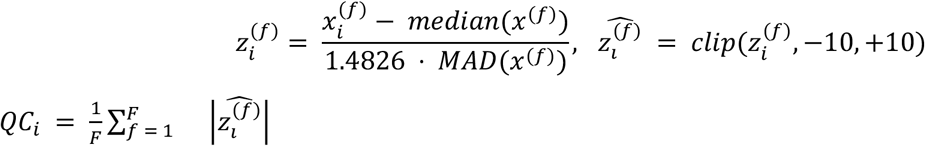

where *F* = 15 (the curated features or principal components) and the factor 1.4826 converts MAD to a consistent estimator of the standard deviation under the Gaussian assumption. Because the analyzed cell populations can be biologically heterogeneous, *QC_i_* was interpreted as a population-relative outlier score rather than a binary classifier of valid versus invalid cells. Within the EpiCycle workflow, labels were assigned to triage tiers based on user-adjusted thresholds on *QC_i_* (and additional features), enabling prioritization of labels for manual review. For exploratory analysis, extracted features were additionally projected into a low-dimensional PCA embedding and displayed interactively alongside the 3D viewer, allowing users to visualize the QC scores over the morphospace for a given stack, lasso-select regions of interest, and highlight corresponding labels in the volume for immediate quality assessment.

#### 7.3 Quality control filtering

Reconstructed cells in the human fetal skin samples were curated for quality in EpiCycle through an expert-guided, human-in-the-loop process. For each stack, 3D cell instances were first surface-smoothed isotropically (σ = 1.5), and the per-cell QC score (§7.2) was computed; higher scores flag cells with unusual size, shape or reconstruction geometry. EpiCycle’s quality control tools allow cells to be triaged in a tiered fashion by dragging thresholds on the QC-score histogram, lasso-selected in the quality-control feature plots (scatter, PCA and UMAP embeddings) and rendered in 3D for inspection **(Fig. S4C)**. Using these tools, poor reconstructions were removed iteratively. Obvious debris fragments, which separated as distinct low-quality populations in the QC-score histogram and UMAP, were removed first; the remaining cells were then manually filtered by cell volume and by nuclear-channel intensity, since a genuine cell contains a nuclear signal that a spurious mask fragment lacks.

In addition to the quality-based triage above, cells at the volume border, whose shapes were truncated by the field of view, were excluded from the morphometric analysis. These cells were only retained as neighbors when constructing the cell contact graph (§7.6), such that the niche of the interior cell was not artificially depleted at the tissue edge.

#### 7.4 Morphospace construction and analysis

For the human fetal morphospace analysis, cell instances aggregated across all retained stacks were characterized using a curated set of 31 morphological features (**Table S3**) spanning cell size, elongation and anisotropy, sphericity and solidity, orientation, skeleton topology, apical-basal cross section asymmetry, and cross-section profile descriptors.

Each feature was preprocessed independently, features with non-negative values and absolute skewness |*γ*_1_| > 1 were logtransformed via *x* ↦ *log*(*1* + *x*) to reduce right-skew; features not meeting the criterion were left untransformed. All features were standardized using `RobustScaler` (median-centered, IQR-scaled) and winsorized at ±5 to limit the influence of extreme outliers. Principal Component Analysis (PCA; via `scikit-learn`) was applied to the preprocessed feature matrix, retaining the first 10 principal components as the morphospace basis for downstream analyses (**Table S6**). Hierarchical clustering of the PC loading matrix was then used to verify that principal axes drew on independent groups of shape descriptors rather than redundant measures of a single feature.

Shape clusters were identified via unsupervised clustering using the leiden algorithm after constructing a k nearest-neighbor (knn) graph within the 10D morphospace. For each cell, the *k* = 15 nearest neighbors were identified using euclidean distance metric; edges were weighted using an exponential distance kernel *w_i_*_j_ = *exp*(−*d_i_*_j_), and duplicate edges from the mutual-neighbor union were combined by taking the mean weight. The resolution parameter was swept over {0.3, 0.5, 1.0, 1.5, 2.0, 2.5, 3.0} using a fixed seed (*seed* = 42) to characterize cluster stability across coarse and fine partitions (**Fig. 6, leiden resolution = 1.0; Video S7**).

#### 7.5 Representative shape sampling

Representative cells were sampled per cluster to provide archetypes for each shape cluster. For each cluster *c*, a reference point *m_c_ ε* ℝ^10^ was defined as the coordinate-wise median of the PC coordinates of its member cells, taken independently over each of the 10 principal components. Cells were then ranked by euclidean distance]|*x_i_* − *m_c_*|]_2_to this reference, computed on unscaled component scores so that components are implicitly weighted by the variance they explain. The single closest cell to the reference point was taken as the cluster archetype, and then the ten closest cells (archetype included) were taken together as a representative gallery capturing the core morphology of each cluster. Cells farther from the reference, which correspond to transitional or outlier shapes near cluster boundaries, were therefore not sampled. The final representative cells were rendered as 3D meshes as illustrated in **Fig. 6D**.

#### 7.6 Graph-diffusion niche embedding

To assess whether morphologically similar cells co-localize spatially, a niche embedding was computed in which each cell’s 10D morphospace coordinates were iteratively blended with those of its physical neighbors over the cell-cell contact graph (constructed in §7.1). The niche embedding was constructed and analyzed per stack: adjacency information was not shared across stacks, and each stack’s adjacency graph was normalized independently.

For each stack, a weighted adjacency matrix 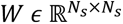 was constructed from EpiCycle-exported contact table, where *W_i_*_j_ is the number of contact voxels shared between cells *i* and *j* (as estimated by graduated binary dilation during EpiCycle feature extraction; §7.1). Contacts with fewer than 100 voxels were filtered out to remove dilation artifacts and structurally weak touches. The adjacency matrix was symmetrized, and row-stochastic (random-walk) normalization was applied:

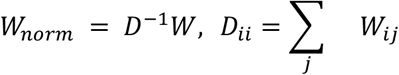

so that each row of *W_norm_* specifies a contact-area-weighted mean over a cell’s direct neighbors. Cells with zero retained contacts (isolated cells after thresholding) were flagged and assigned 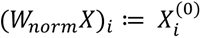 at every iteration to preserve their original morphospace coordinates.

Let 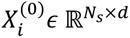 denote the original PC coordinates (with d = 10) for the cells in a given stack. The niche embedding is computed by repeatedly blending each cell’s intrinsic morphology with a contact-weighted average of the cells it physically touches:

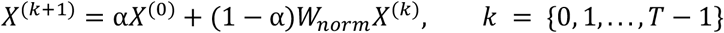

where *a ε* [0, 1] is self-weight, anchoring the output toward each cell’s original morphology, *T* is the number of iterations, and the final niche embedding *<u>X</u>* = *X*^(*T*)^is the per-cell output. In practical terms *a* controls how strongly a cell retains its own original shape information versus borrowing information from its physical neighbors, whereas *T* controls the effective spatial range over which neighbor information can spread through the contact graph: *T* = 1 uses direct neighbors only *T* = 2 incorporates neighbors-of-neighbors, and larger *T* values effectively capture progressively broader tissue context with each iteration allowing information to diffuse further. Setting *a* = 0 yields a pure weighted mean of neighbors while *a* = 1 returns the original PCs unchanged. **Fig. 6F** illustrates this update for a single cell with ⍺ = 0, where P_ij denotes the entry (W_norm)_ij.

Per-stack niche matrices were reassembled into a single global niche matrix of shape (*N*, *d*); cells from stacks lacking adjacency data were flagged as invalid and excluded from downstream analyses. leiden clustering was applied at resolutions {0.3, 0.5, 1.0, 1.5, 2.0, 2.5, 3.0}, and UMAP was used to visualize the niche space (`n_neighbors = 15`, `min_dist = 0.1`). Results shown in **Fig. 6** utilize ⍺ = 0, T = 3 and leiden resolution 1.5 on representative stacks; **Video S7** shows configurations spanning one to five iterations and ⍺ ∈ {0.0, 0.3, 0.5}.

#### 7.7 Hair placode and fingerprint ridge comparisons

Groups were defined within the front trunk and ventral digit samples by niche cluster (resolution 1.5): hair placode, N17; its surroundings, N0; ridge basal layer, N3, N7, N10, N13 and N14; ridge surroundings, all other cells of the ventral digit sample. Each structure was compared with the surroundings from its own sample. Cell orientation is the angle between a cell’s major axis and the imaging z axis, which is approximately perpendicular to the skin surface. Distances between groups (**Fig. 7M**) are the distance between group medians in the 10D morphospace, divided by the mean of the two groups’ spreads, where a group’s spread is the median distance of its cells to the group median. Representative cells (**Fig. 7H**) are the ten cells of each group closest to the group median in the 10D morphospace sampled in the same manner in §7.5.

Differences among the four groups **(Fig. 7I–L)** were tested per feature with Welch’s one-way ANOVA followed by Games–Howell post hoc comparisons. Cell volume was log-transformed before testing. Cells were treated as replicates; all four groups come from a single donor.

## ACKNOWLEDGEMENTS

We thank the Life Science Building Aquatics staff for animal care and the Tillicum team at the University of Washington for computing support. The authors are grateful to Silvia Fre for comments on the manuscript, Laure Bally-Cuif and Nicolas Dray for discussion, support and encouragement, and all members of the Rasmussen lab for feedback and inspiration on the project.

## AUTHOR CONTRIBUTIONS

Conceptualization: E.J.A.Q; Data Curation: E.J.A.Q.; Formal Analysis: E.J.A.Q.; Funding Acquisition: E.C.B., D.J.H., J.P.R.; Investigation: E.J.A.Q., E.C.B., E.P.; Methodology: E.J.A.Q.; Project Administration: E.J.A.Q., J.P.R.; Resources: L.H., D.J.H., J.D.G.; Software: E.J.A.Q.; Supervision: J.P.R.; Validation: E.J.A.Q., J.P.R.; Visualization: E.J.A.Q.; Writing – Original Draft Preparation: E.J.A.Q., J.P.R.; Writing – Review & Editing: E.J.A.Q., E.C.B., E.P., L.H., D.J.H., J.D.G., J.P.R.

## DATA AND RESOURCE AVAILABILITY

EpiCycle is open-source software available at https://github.com/eL-Gene/EpiCycle. Confocal image volumes for the zebrafish, mouse salivary gland, human fetal skin datasets (Table S1) and associated data for analyses featured in this study are deposited at BioImage Archive [accession S-BIAD4434]. The analysis notebooks that generate the benchmark values (Table S2), the morphospace and niche analyses, and all plots in Figures 2, 3, 6, 7, S2 and S3 are available at https://github.com/eL-Gene/EpiCycle-analyses.

## DECLARATION OF GENERATIVE AI AND AI-ASSISTED TECHNOLOGIES

Large-language models were selectively used during the development of EpiCycle software. The authors used Claude Code models (Opus 4.6-5.5, by Anthropic) to assist with writing and refactoring code for the graphical user interface, testing, documentation and code optimization. All code was reviewed and tested by the authors. All code pertaining to experimental design, the analysis presented in this work and conceptualization of the workflow was generated by the authors. The authors take full responsibility for the content of the publication.

## Supplementary Video Legends

**Video S1. Tissue-scale 3D instance segmentations of adult zebrafish trunk epidermis with domain-specialized models, related to Fig. 4A**

3D rendering of a tiled confocal volume of adult zebrafish trunk epidermis, showing E-cadherin (green; *TgBAC(cdh1:cdh1-sfGFP)*) and nuclei (magenta; *Tg(actb2:h2b-2x-mScarlet)*). The lateral and axial domain-specialized model pair was applied across the whole volume using the tiled reconstruction pipeline (1024 × 1024 × Z tiles, 25% overlap) without further training. Each color marks one reconstructed cell. The video shows the raw volume, the full reconstruction, close-ups of representative basal keratinocytes (compact, polygonal, at the dermal side) and superficial keratinocytes (flattened and anisotropic), and finally examples of poor reconstructions (e.g., axial merges, rough surfaces and incoherent instances), which concentrate where membrane signal is weak or in deeper tissue regions.

**Video S2. Domain-specialized models applied to wounded adult zebrafish epidermis, related to Fig. 4B**

3D rendering of wounded adult zebrafish scale epidermis expressing reporters for E-cadherin (green; *TgBAC(cdh1:cdh1-sfGFP)*) and nuclei (magenta; *Tg(actb2:h2b-2x-mScarlet)*), reconstructed with the adult domain-specialized model without retraining. Each color marks one cell. The wound and the surrounding unwounded epidermis are shown.

**Video S3. Targeted fine-tuning of domain-specialized model on wounded adult zebrafish epidermis, related to Fig. 4C**

A 1024 × 1024 × Z subregion of the wounded adult zebrafish epidermis (E-cadherin, green; nuclei, magenta) shown as the curated 3D ground truth (left), the reconstruction from the original domain-specialized model (top right) and the reconstruction after targeted fine-tuning (bottom right). The initial domain-specialized model reconstruction was corrected in EpiCycle’s multiview annotator; lateral and axial models were then fine-tuned on 2D slices from four local sub-stacks only (Fig. 4C5), with the rest of the annotated area withheld. The fine-tuned pair was applied to the full tile. The volume is shown from above, from the side and in cross-section with a moving clipping plane so that labels can be compared with the membrane signal. Instance separation and boundary quality improve for the previously undersegmented morphologies, including in the withheld and adjacent regions. Cell colors are assigned independently in each panel.

**Video S4. Domain-specialized models applied to developing zebrafish epidermis, related to Fig. 4D**

3D reconstructions of juvenile zebrafish trunk epidermis at four stages of scale formation: pre- placode, placode, scale condensate and maturing scale. Fish carried the same E-cadherin (green) and nuclear (magenta) reporters as the adult training data (shown in Videos S1–S3, Figs. S1 and 2–4). Each stack was segmented with the adult domain-specialized model pair without further training and fused in 3D with u-Segment3D. The raw signal is shown first, then the reconstructed cells (one color per cell) and then edge-on views with cell outlines over the membrane signal. Basal cells are resolved at all stages; separation failures concentrate along the axial direction where the membrane signal between vertically adjacent cells is discontinuous and decrease as lateral boundaries sharpen in later stages. Only the epidermis was segmented.

**Video S5. 3D cell and nucleus reconstructions of an embryonic mouse salivary gland explant, related to Fig. 5A1-2,B1-2**

Two subvolumes of an E14.5 mouse salivary gland explant cultured for 48 h and stained for E-cadherin (green) and DAPI (magenta): an end-bud region (top row; stack shown in Fig. 5A1) and a ductal **branch** region (bottom row; stack shown in Fig. 5A2). Cells (left column) and nuclei (right column) were reconstructed with separate lateral and axial model pairs trained from scratch in EpiCycle.

**Video S6. 3D cell reconstructions and morphological domains of human fetal skin, related to Figs. 5C–F and 6H**

Four of the 13 reconstructed human fetal skin stacks (12–14 weeks EGA; α-catenin, green; DAPI, magenta): unpatterned epidermis, a hair placode (front trunk), fingerprint ridges (ventral digit) and a fingerprint whorl (ventral digit), corresponding to Fig. 5C–F. Each stack is shown as a raw signal, then as the 3D cell masks produced by the human-adapted model pair after quality control and then with cells colored by shape cluster (leiden resolution 1.0 on the 10-PC morphospace; Fig. 6D) and by niche cluster (leiden resolution 1.5 on the graph-diffused niche embedding, α = 0, T = 3; Fig. 6G). Niche clusters are then isolated one at a time, with the remaining cells rendered translucent.

**Video S7. Shape and niche clusters across leiden resolution and niche-diffusion parameters, related to Fig. 6H**

The four human fetal skin stacks of Video S6, rendered in 3D. Cells are first colored by shape clusters for leiden resolutions 0.3, 0.5, 1.0, 1.5, 2.0, 2.5 and 3.0 on the 10D morphospace (Fig. 6 uses resolution = 1.0). Cells are then colored by niche clusters for one to five diffusion steps (T) at self-weight α = 0, 0.3 and 0.5, with leiden resolution fixed at 1.5 (Fig. 6 uses α = 0, T = 3). Parameter values are shown above each panel. Cluster colors are assigned independently at each setting. The placode and ridge domains persist across settings, while increasing α anchors each cell toward its own shape and progressively blurs the domains.

## Supplementary Tables

**Table S1. Datasets, imaging and preprocessing parameters.**

**Table S2. Benchmark values underlying Figures 2, 3, S2 and S3: 2D AP@50 per HITL round and 3D AP@50 per stack and training condition.**

**Table S3. Feature dictionary.**

**Table S4. Reagents and software.**

**Table S5. Model training data and hyperparameters.**

**Table S6. Segmentation, reconstruction and analysis parameters by dataset.**

## SUPPLEMENTARY INFORMATION

### Supplementary Figures and Figure Legends

**Figure S1.**
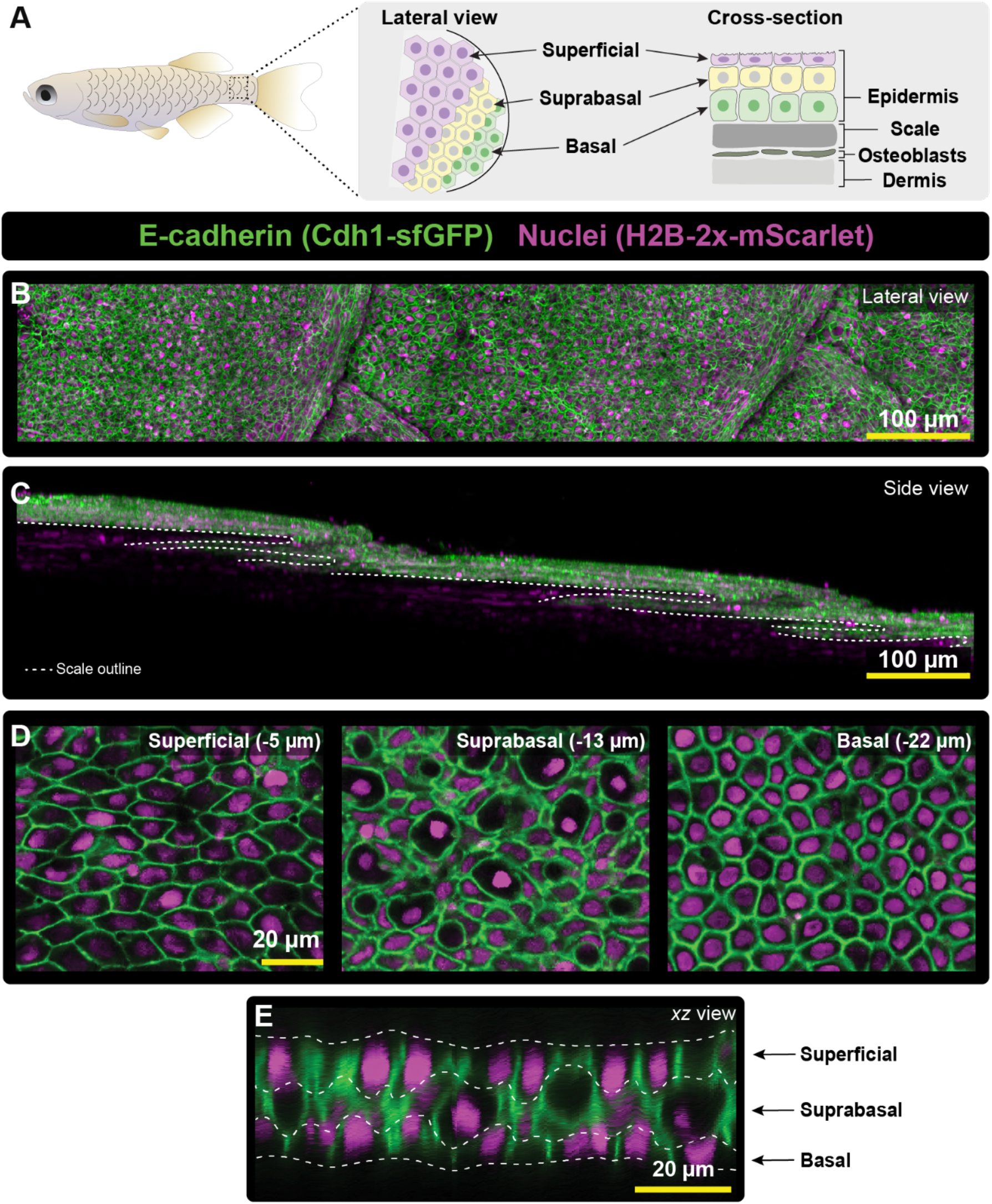
Zebrafish trunk epidermis as a model stratified epithelium. **(A)** Schematic of adult zebrafish skin anatomy illustrating a stratified epidermis overlying a scale. The trunk epidermis contains three distinct layers: superficial (purple), suprabasal (yellow) and basal (green). **(B)** Representative tile-scan confocal image of adult trunk epidermis from a *TgBAC(cdh1:cdh1-sfGFP);Tg(actb2:h2b-2x-mScarlet)* fish, showing E-cadherin/Cdh1-sfGFP at cell-cell junctions (green) and H2B-2x-mScarlet in nuclei (magenta). Scales appear as repeating structures across the tissue. **(C)** Corresponding side view as in **(B)** showing epidermis overlying and wrapping around the posterior margin of scales (outlines). **(D)** Single optical sections at progressive imaging depths through the trunk epidermis revealing layer-specific morphologies. **(E)** Orthogonal *xz* optical section through the trunk epidermis illustrating stratified organization of superficial, suprabasal and basal layers along tissue depth axis. Dashed contours indicating approximate corresponding layer boundaries shown in (A).

**Figure S2.**
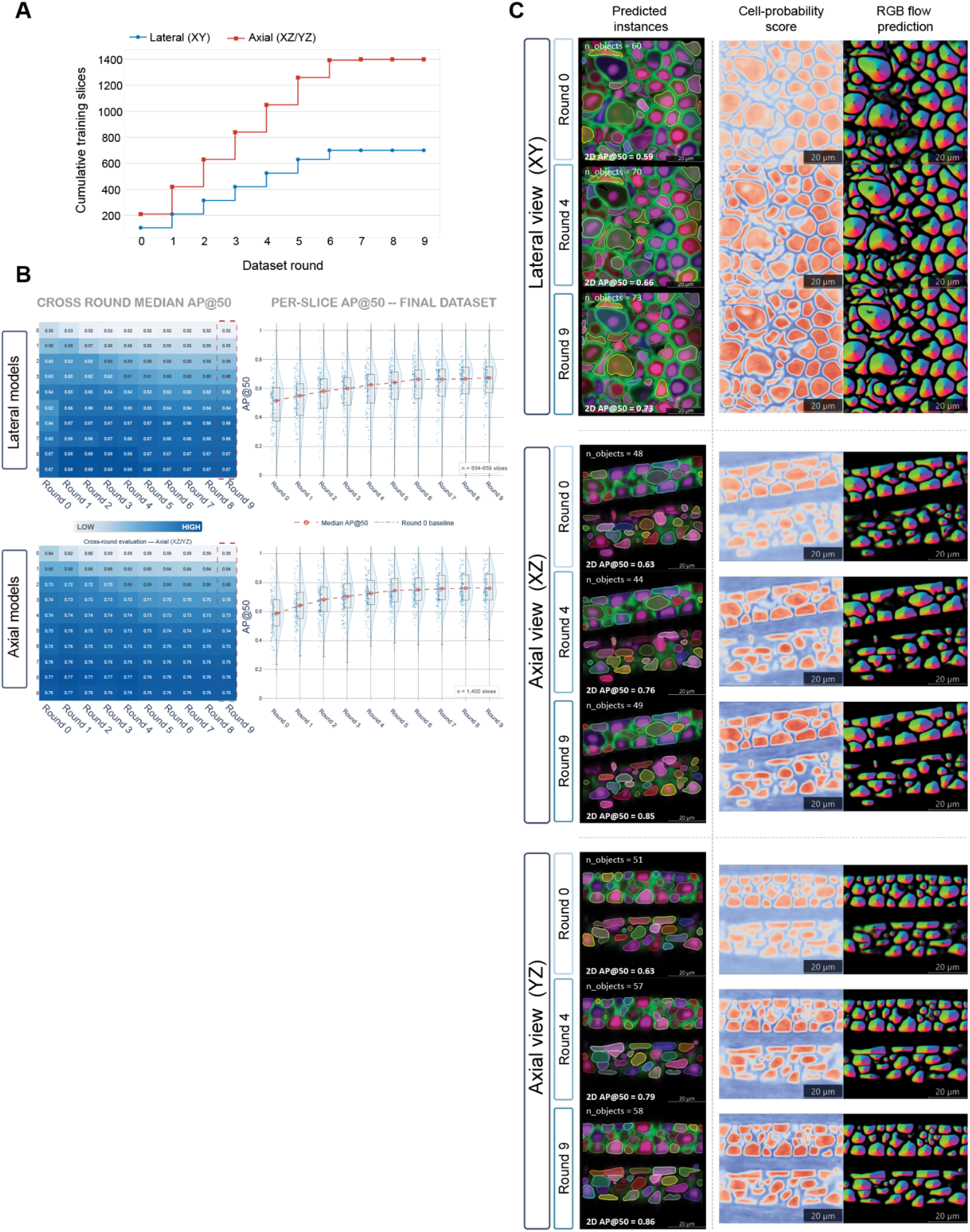
Human-in-the loop training of lateral and axial 2D models across rounds. **(A)** Cumulative number of 2D training slices at each dataset round for lateral (XY; blue) and axial (XZ/YZ; red) models. Each round added 15 lateral and 30 axial slices per development stack with training set saturation at 700 lateral and 1,400 axial slices by round 6. **(B)** Left: complete cross-round median 2D AP@50 for lateral (top) and axial (bottom) models across all ten rounds. Box plot shows median and interquartile range, with jittered points and kernel density overlaid. Values in **Table S2**. **(C)** Representative lateral and axial slices segmented by models from rounds 0, 4, 9 (as shown in Figure 2). Columns show predicted instances derived from Cellpose cell-probability map, and the predicted flow field (RGB-encoded direction) using default cell probability and flow threshold parameters (see Methods).

**Figure S3.**
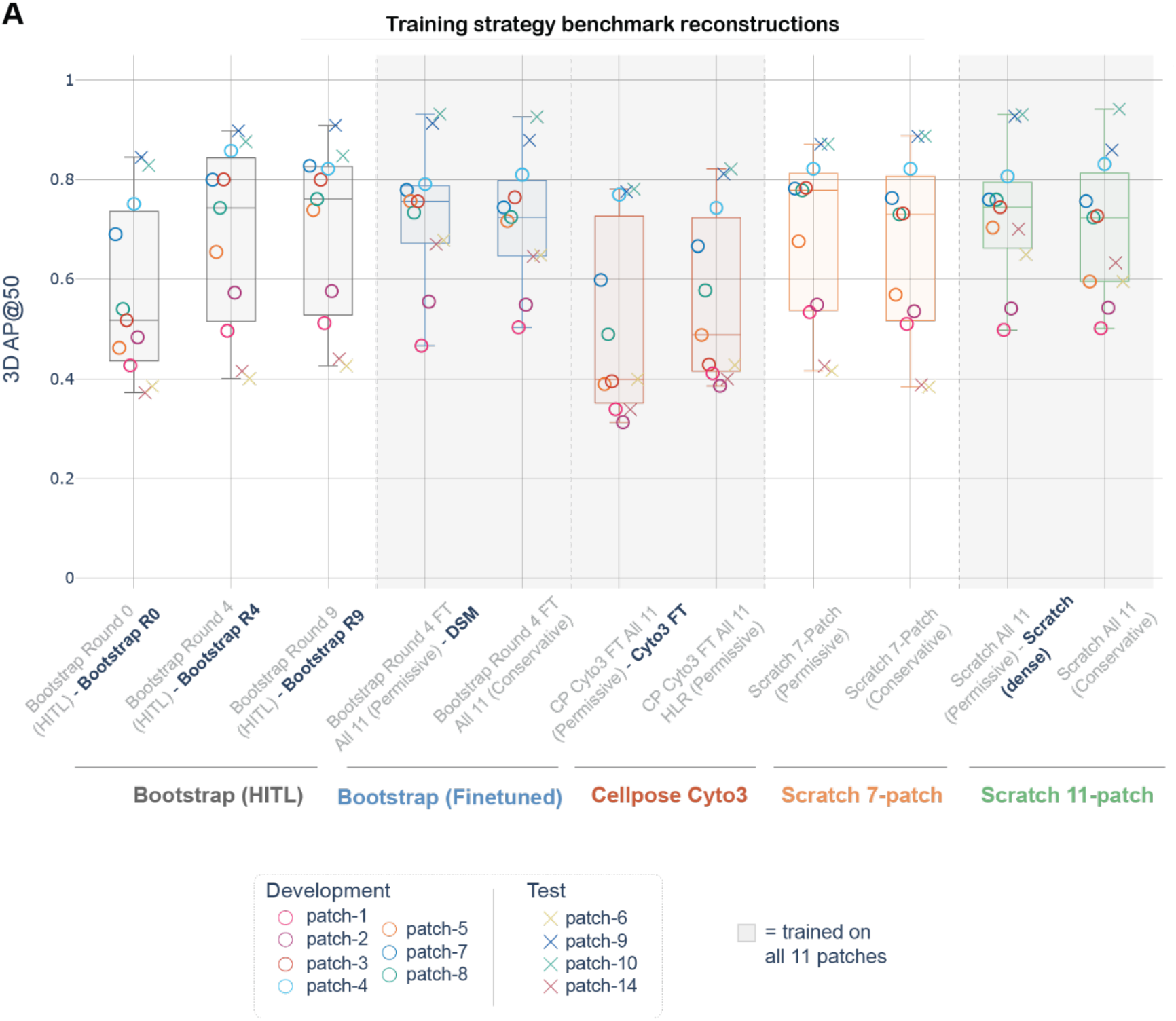
3D reconstruction accuracy across all eleven training configurations. **(A)** Complete per-stack 3D AP@50 for every training condition, each evaluated on 11 ground-truth stacks. Box plot showing median and interquartile range (as shown in Figure 3). Conditions, left to right: Bootstrap (2D HITL) rounds 0, 4, and 9; Bootstrap round 4 on all 11 stacks with data permissive (DSM model) or conservative training/validation/test splits; Cellpose cyto3 fine-tuned on all 11 stacks at learning rates 0.01 (Cyto3FT) or 0.1 (high learning-rate, HLR); models trained from scratch on 7 development stacks, under data permissive or conservative regimes; and models trained from scratch on all 11 stacks under permissive (Scratch, dense) or conservative regimes. Conditions named in bold are those shown in Figure 3C and 3E. Per-stack values in **Table S2**.

**Figure S4.**
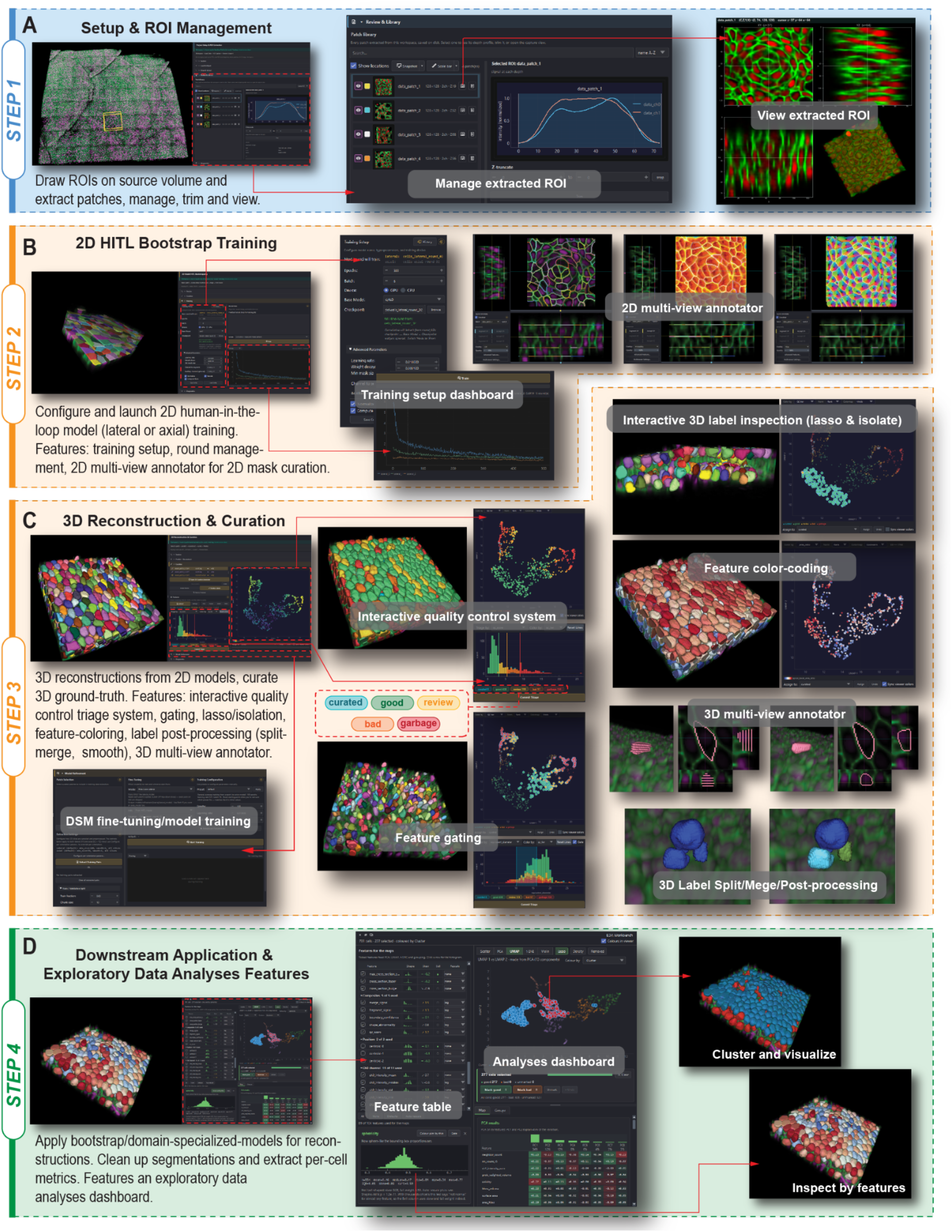
The EpiCycle plugin provides an integrated interface for 2D HITL training, 3D reconstruction and curation and downstream exploratory data analysis. **(A)** *STEP1 Setup and ROI management*. Regions of interest (ROIs) are selected from a loaded volume and extracted as patches and managed accordingly through the library manager section allowing users to trim extracted patches and visualize individual patches accordingly. Additional features such data conversion (not shown) and initial optional preprocessing included (not shown) are also included. **(B)** *STEP2 2D HITL bootstrap training*. Training parameters for lateral and axial models are configured through the training interface, with training progress visualized in real-time. Predictions and additional labels added within each round of training can be reviewed and corrected in synchronized multi-view annotators displaying segmentation masks and associated Cellpose outputs. **(C)** *STEP3 3D reconstruction and curation*. Predictions from 2D models are reconstructed into 3D instances using u-Segment3D and inspected through an interactive quality-control workflow. Curation tools include feature-based visualization and gating, lasso-based cell isolation, multiview-label editing, slice interpolation, label splitting and merging, and post-processing. Curated annotations can additionally be used to fine-tune or train domain specialized models. **(D)** *STEP4 Downstream application and exploratory analyses*. Trained models can be applied to larger ROIs. Per-cell features can subsequently be extracted and examined in EpiCycle exploratory analysis workbench, which supports feature tables, distributions, low dimensional embeddings (PCA/UMAP/t-SNE), clustering, and interactive visualization of resulting cells in the same manner shown in *STEP3*.

